# Yellow adipocytes comprise a new adipocyte sub-type present in human bone marrow

**DOI:** 10.1101/641886

**Authors:** Camille Attané, David Estève, Karima Chaoui, Jason Iacovoni, Jill Corre, Mohamed Moutahir, Philippe Valet, Odile Schiltz, Nicolas Reina, Catherine Muller

## Abstract

During energy demanding conditions, white adipocytes store triglycerides and release fatty acids through lipolysis. In contrast, bone marrow adipocytes (BM-Ad) increase in size during caloric restriction, suggesting this fat depot exhibits precise metabolic specificity. We found subcutaneous adipocytes (SC-Ad) and BM-Ad share morphological features, but possess distinct lipid metabolism. BM-Ad show enrichment in cholesterol-oriented metabolism that correlates with increased free cholesterol content, while proteins involved in lipolysis were downregulated. A strong down-regulation in expression of monoacylglycerol (MG) lipase was observed leading to an accumulation of major MG species and accordingly the basal and induced lipolytic responses were absent in BM-Ad. These features are not recapitulated *in vitro* using differentiated bone marrow mesenchymal stem cells. Since our data demonstrate that BM-Ad comprise a distinct class of adipocytes, we propose renaming them yellow adipocytes.

## Introduction

In mammals, white adipose tissue (WAT) accumulates at various sites throughout the body. The most important and well-studied fat deposits occur in subcutaneous regions (SC-AT) and in the abdominal cavity surrounding key internal organs like the pancreas and intestines (Zwick et al, 2018). Other adipose-specific deposits also form around the heart, kidney, prostate in men and mammary glands in women (Zwick et al, 2018). In addition to WAT, mammals also possess brown adipose tissue (BAT) located in the interscapular and supraclavicular regions, representing less than 5% of the total fat mass (Leitner et al, 2017; Nedergaard et al, 2007; Saito et al, 2009; Zingaretti et al, 2009). Brown adipocytes participate in non-shivering thermogenesis and possess a specific morphology that includes several small lipid droplets and high mitochondria content (Bartelt & Heeren, 2014; Cinti, 2001). In contrast, white adipocytes store energy as triglycerides (TG) in their unique large lipid droplet (LD) after energy intake and release free fatty acids (FFA) through lipolysis in energy demanding conditions (Zechner, 2015). Lipolysis occurs through a biochemical pathway that uses consecutive actions of adipose triglyceride lipase (ATGL), which catalyzes the conversion of TG to diacylglycerols (DG) and hormone-sensitive lipase (HSL) and hydrolyzes DAG to monoacylglycerols (MG), monoacylglycerol lipase (MAGL) and the newly identified α/β hydrolase domain-containing protein 6 (ABHD6), which hydrolyzes MG to FA (Zhao et al, 2016) and glycerol (Zechner, 2015). White adipocytes also have an important endocrine function as they can release multiple soluble factors called adipokines, such as leptin and adiponectin (Fasshauer & Blüher, 2015).

One intriguing adipose tissue (AT) localizes to the bone marrow called bone marrow adipose tissue (BM-AT) that constitutes over 10% of the total fat mass in lean and healthy humans (Cawthorn et al, 2014). Technological advances in quantitative imaging of BM-AT in both mice and humans revealed that BM-AT presents unique features that highlight their physiological specificity. Many studies demonstrated that BM-AT increases in different pathophysiological conditions such as aging (Justesen et al, 2001; Scheller et al, 2015), osteoporosis (Justesen et al, 2001; Yeung et al, 2005) and obesity (Bredella et al, 2010; Doucette et al, 2015). These findings suggest this adipocyte population plays a larger role beyond that of “filler-cells”. In stark contrast to the other WAT, the number and size of bone marrow adipocytes (BM-Ad) also increase during caloric restriction conditions in mice (Cawthorn et al, 2014; Devlin et al, 2010), rabbits (Bathija et al, 1979; Tavassoli, 1974) and human patients suffering from anorexia nervosa (Abella et al, 2002; Bredella et al, 2010). Decreases in bone marrow adiposity occurs only in severe nutrient deprivation in rabbits (Cawthorn et al, 2016) and late stages of anorexia nervosa associated with gelatinous transformation of the bone marrow (BM) [(Abella et al, 2002); for review (Ghali et al, 2016)].

Given the significant role for AT in regulating energy homeostasis, it is critical to elucidate why this tissue copes with changes in energy status in such a specific way that leads to still store and not dispense fuel when needed. However, knowledge of the phenotype of primary BM-Ad in physiology is sparse and hampered by difficulty to obtain sufficient isolated BM-Ad in mice and from harvesting human BM-AT partly due to the physical location (inside bone). Most studies on BM-Ad use rodents or human *in vitro* models. Mouse studies indicate BM-Ad regulate hematopoiesis and bone mass (Naveiras et al, 2009; Zhou et al, 2017). However, species-specific differences between rodent and human BM-AT exist that reinforce using caution when extrapolating information across species (Scheller et al, 2016). Two types of adipocytes in mouse have been described: regulatory and constitutive BM-Ad (rBM-Ad and cBM-Ad, respectively) (Scheller et al, 2015). cBM-Ad are present in tail vertebrae and the medullary canal from the tibia-fibular junction into the malleolus. However, rBM-Ad develop postnatally within the BM of long bones extending from below the growth plate through the metaphysis and into the diaphysis (Scheller & Rosen, 2014). Existence of these two populations remains unconfirmed in humans. Inside the diaphysis of long bone, the number of BM-Ad varies between mouse strains and species, and some strains require pharmacological induction of BM-Ad by drugs such as glucocorticoids and thiazolidinedione (Scheller et al, 2016). Yet, human BM-Ad consistently fill 50 to 70% of the bone marrow cavity (Hindorf et al, 2010). Many studies use bone marrow mesenchymal stromal cells (BM-MSC) differentiated in adipocytes *in vitro*. However, it is unclear whether these differentiated cells recapitulate the phenotype of mature human primary BM-Ad. These *in vitro* studies suggest a role for BM-Ad in hematopoiesis regulation (Mattiucci et al, 2018; Naveiras et al, 2009), bone remodeling (Hardaway et al, 2015) and cancer progression (Diedrich et al, 2016; Herroon et al, 2013; Liu et al, 2015; Shafat et al, 2017; Tabe et al, 2017). These issues highlight that our knowledge of the physiological phenotype of primary BM-Ad remains limited. Using combined lipidomic and proteomic large-scale approaches, we purified and characterized human BM-Ad harvested from the femoral diaphysis of patients undergoing hip surgery with paired subcutaneous adipocytes (SC-Ad) and found that BM-Ad exhibit clearly distinct lipid metabolic features that reveal a new adipocyte sub-type.

## Results/Discussion

### Isolated SC-Ad and BM-Ad share morphological properties of white adipocytes

After harvesting paired SC-AT and BM-AT from patients undergoing hip replacement surgery, we isolated adipocytes after collagenase digestion (Fig 1A). In AT from both locations, the vast majority of the space contained large and cohesive mature adipocytes with a unique LD filled with neutral lipids (assessed by Bodipy staining) (Fig 1B). Mature adipocytes from both locations expressed perilipin 1 (PLIN1) at the surface of the LD (Fig EV1A-B) and exhibited a very thin cytoplasm rim, a morphological trait expressed by white adipocyte (Fig EV1B) (Cinti, 2001). SC-AT and BM-AT also contained blood vessels highly positive for actin staining and stroma vascular cells at both stromal and perivascular positions (Fig 1B and Fig EV1A). Using transmission electron microscopy approach, we observed that both SC-Ad and BM-Ad present in the AT display a large LD surrounded by a very thin cytoplasm with the nucleus located at the cell periphery between the plasma membrane and the LD (Fig 1C). We performed an enzymatically based digestion protocol to isolate adipocytes from both tissues. After obtaining a population of cells constituted only of adipocytes, our results indicated that our tissue dissociation preserved the morphological identity of the isolated adipocytes. The isolated BM-Ad and SC-Ad shared the same morphology found within the tissues characterized by the presence of a unique and large LD filled with neutral lipids (Fig 1D). In addition, F-Actin staining showed a similar cytoskeleton architecture between the two types of cells (Fig 1D). Taken together, our results demonstrate the BM-AT present in the diaphysis of long bone is composed of cohesive adipocytes that exhibit the morphological appearance of white adipocytes as assessed by their unique LD, surrounded by a thin cytoplasm and a nucleus present at the periphery of the cells. With the caution noted in the introduction, a recent study in mice that used electronic transmission found that BM-Ad exhibit similar rounded morphology with a unique large LD (Robles et al, 2019). Yet, a recent report suggested that mouse BM-Ad express some genes related to BAT, including PRDM16 and FOXC2 (Krings et al, 2012). However, this study used whole tibia extracts, which contain adipocytes and contaminating cells, including myeloid cells and osteoblasts that express PRDM16 and FOXC2, respectively (Kim et al, 2009; Nishikata et al, 2011). Here, we present an initial morphological characterization of human BM-Ad, where they exhibit traits of white adipocytes that do not clearly distinguish them from “classical” white adipocytes.

**Fig 1:**
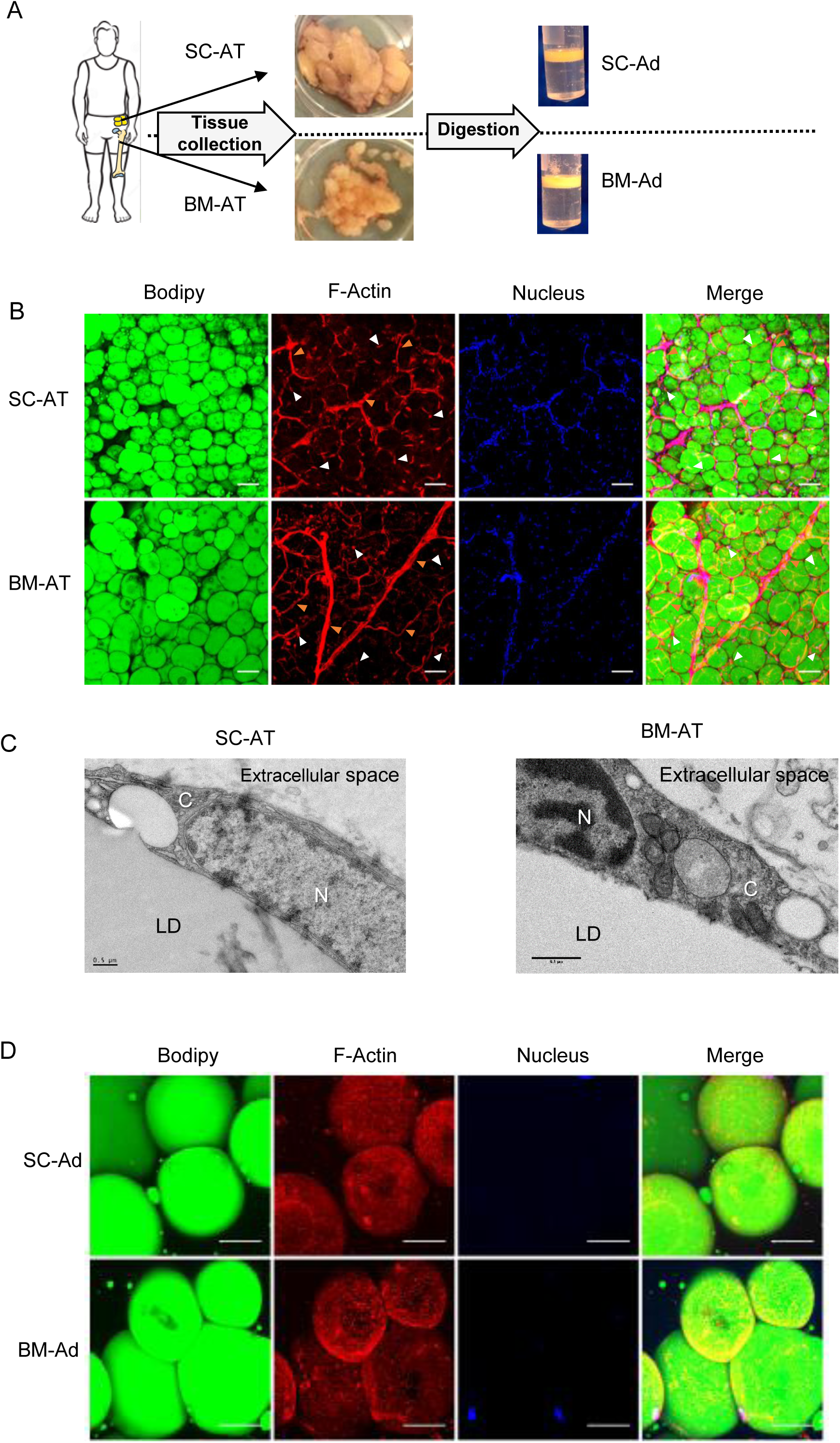
SC-Ad and BM-Ad exhibit similar morphological properties. **A.** Scheme of the experimental protocol designed to obtain paired human primary bone marrow (BM-Ad) and subcutaneous adipocytes (SC-Ad). Paired bone marrow (BM-AT) and subcutaneous adipose tissues (SC-AT) were harvested from patients undergoing hip replacement surgery. BM-AT, that share the same macroscopic aspects compared to SC-AT, was isolated from the red bone marrow containing hematopoietic cells. After enzymatic digestion, floating cells were rinsed and collected for subsequent experiments. **B.** Whole mount SC-AT and BM-AT were stained with Bodipy 493/503 (neutral lipids, green), Phalloidin (F-Actin, red) and TOPRO-3 (nucleus, blue). Z stack images were taken using confocal microscope with 10X objective (n=3). Representative maximum intensity projection is shown. Orange arrowheads show vessels. White arrowheads show stromal cells. Scale bar, 100µm. **C**. Transmission electron microscopy images of SC-AT and BM-AT. N: Nucleus, LD: Lipid Droplet, C: cytoplasm. Scale bar, 0.5µm. **D**. Primary SC-Ad and BM-Ad were isolated and stained with Bodipy 493/503 (neutral lipids, green), phalloidin (F-actin, red) and TOPRO3 (nucleus, blue). Z stack images were taken using confocal microscope with 40X objective (n=3). Representative maximum intensity projection is shown. Scale bar, 50µm.

### Lipid profile in BM-Ad reveals enriched diverse lipid species like monoacylglycerol (MG) and cholesterol

We then further characterized the phenotype of BM-Ad by studying their lipid profile compared to SC-Ad. Lipids were extracted from tissues and isolated adipocytes. As shown in Fig 2A, each tissue and isolated adipocytes showed a similar total lipid content. A quantitative LC-MS/MS based analysis of the total lipid content extracted from BM-Ad and SC-Ad was performed using a recently developed approach that uses both positive and negative ionization modes to cover the largest spectrum of detectable lipid species (Breitkopf et al, 2017). The analysis structurally characterized and identified 818 lipid species originating from the main lipid categories that belonging to 15 lipid classes. The majority of identified lipid species were glycerolipids (GL), including triacylglycerol (TG, 95 %) and diacylglycerol (DG, 2.1%). Beyond GL, the remaining lipids contain a large spectrum of phospholipids (PL), in particular phosphatidylcholine (PC, 1.9%), a major membrane constituent (Wen et al, 2018), sphingolipids (SL) and fatty esters (Fig 2B).

**Fig 2:**
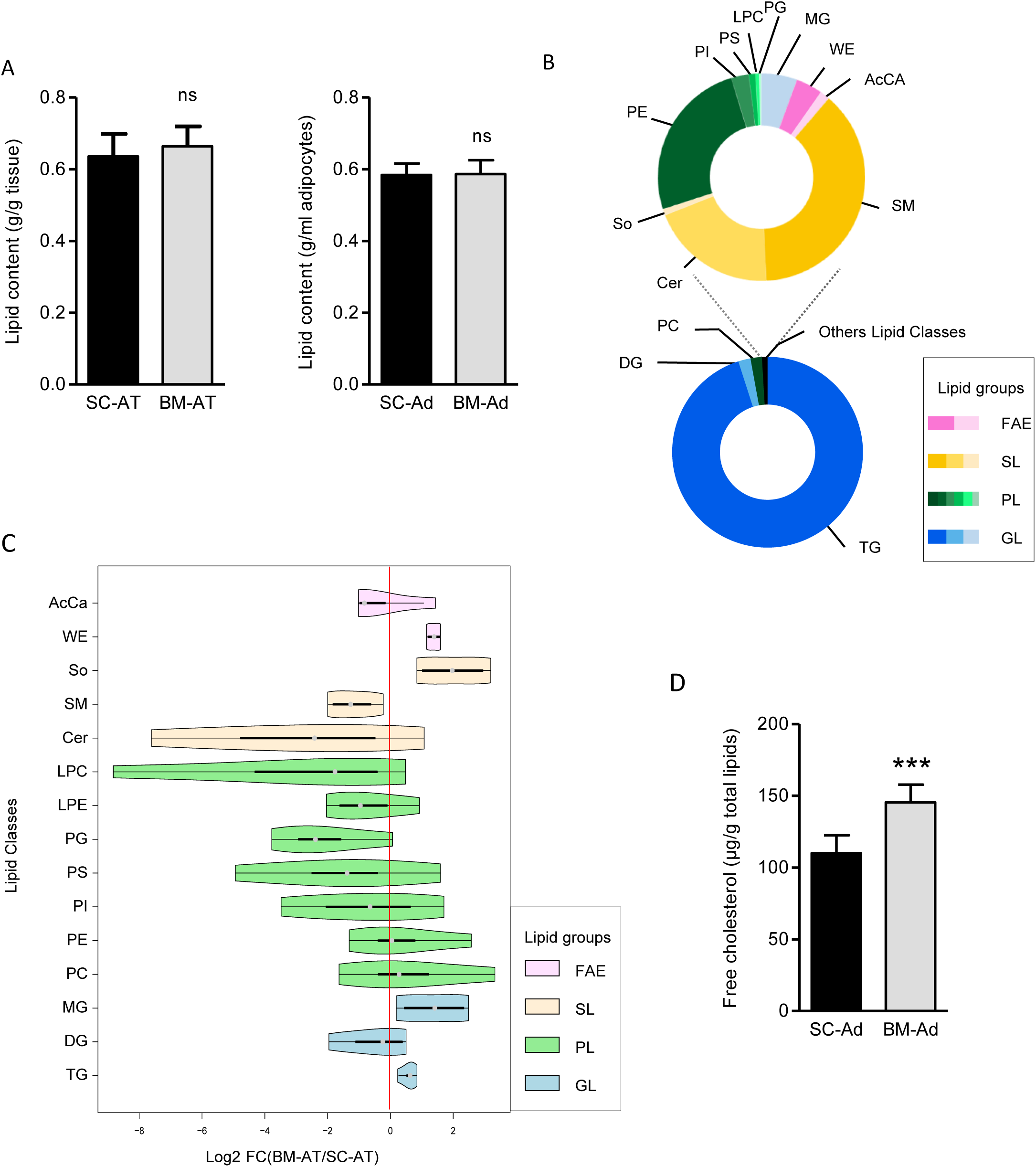
Detailed lipid species analysis of BM-Ad shows increase free cholesterol and MG contents compared to SC-Ad. **A.** Total lipid content in SC-AT and BM-AT (left, n=7) and in SC-Ad and BM-Ad (right, n=17) were extracted and weighted. The quantity of lipids was normalized to the quantity of tissue or the volume of adipocyte from which lipids were extracted. Histograms represent mean ± SEM. ns stands for non-significant according to paired t test. **B.** Pie chart of the relative abundance of the detected lipid classes using large-scale LC-MS/MS approaches. The glycerolipids (GL) are shown in blue shades. TG: triacylglycerol, DG: diacylglycerol, MG: monoacylglycerol; Phospholipids (PL) are in green shades. PC: phosphatidylcholine, PE: Phosphatidylethanolamine, PG: phosphatidylglycerol, PI: Phostatidylinositol, PS: Phosphatidylserine, LPC: Lysophosphatidylcholine, LPE: Lysophosphatidylethanolamine, Sphingolipids (SL) are in yellow shades. SM: Sphingomyeline, Cer: Ceramides, So: Sphingosine; and fatty acid esters (FAE) are in pink shades. WE: Wax Ester, AcCA: Acyl Carnitine. **C**. Violin plot representing the log2 fold change of the 15 lipid classes identified in BM-Ad compared to SC-Ad analyzed by LC-MS/MS (n= 4). The quantity of lipid classes were calculated as the sum of the different lipid species belonging to the same classes. **D**: Free cholesterol contents were measured using an enzymatic assay on lipid extracted from BM-Ad and SC-Ad. The results were normalized to the quantity of total lipids. The histograms represent mean ± SEM, *** p<0. 001 according to paired t test (n= 11).

Unsupervised multivariate analyses of our lipidomic dataset indicated the variance between samples predominantly arose through inter-individual variability (Fig EV2A-B) indicating the lipids stored within mature adipocytes likely came from food intake. We conclude that BM-Ad lipid composition reflects dietary lipid intake, which is consistent with a prior report in SC-Ad (Hodson et al, 2008). We then investigated if differences exist in lipid classes between SC-Ad and BM-Ad by comparing all quantified lipid species for one class between the two locations. As shown in Fig 2C, we observed differences in GL content. BM-Ad, in all samples, exhibited a slight increase in TG content with no changes in DG levels. MG content increased (Fig 2C), which reflected an increase of both saturated and unsaturated major MG species (Fig EV2C). These results suggest that the hydrolysis of MG is not efficient in BM-Ad. Two additional lipid classes are also increased in BM-Ad compared to SC-Ad, wax esters and sphingosine. Of note, only three sphingosine species were detected. The LC-MS/MS approach we used to quantify the lipid species does not identify cholesterol species, a key lipid species contained in adipocyte LD (Schreibman & Dell, 1975). Using a colorimetric assay, we found that BM-Ad showed a 1.5-fold increase in free cholesterol content compared to SC-Ad (Fig 2D). Cholesterol ester was not detected in either sample as we predicted since the vast majority of cholesterol is expressed in a free form in adipocytes (Schreibman & Dell, 1975). Here, we characterize the lipid content in BM-Ad using unsupervised lipidomic approaches. Inter-individual variability suggests that BM-Ad lipid content partially reflects dietary intake as in other adipose depots. Our results demonstrate that intrinsic differences exist between BM-Ad and SC-Ad regarding free cholesterol and MG contents.

### Proteome of BM-Ad and SC-Ad differentiates adipocytes in lipid metabolic functions

We sought to further decipher the metabolic pathways specifically present in BM-Ad, so we conducted a large-scale proteomic analysis on paired SC-Ad and BM-Ad. We detail the data analysis general strategy in Fig EV3A. After data quality control, 3259 proteins were robustly detected. Interestingly, when we searched for proteins known to be secreted by adipocytes, termed adipokines (Fasshauer & Blüher, 2015), our dataset did not highlight significant differences between the two types of adipocytes and K-means clustering based on the expression pattern of adipokines did not allow clustering of the samples according to their anatomical location (Fig EV3B). BM-Ad expressed the adipocyte-specific adipokine, adiponectin, at the same levels as SC-Ad (Fasshauer & Blüher, 2015) (ADIPOQ, Table EV1). We obtained similar results for leptin (LEP, Table EV1), a hormone predominantly produced by adipocytes (Fasshauer & Blüher, 2015). These results indirectly assess the quality of cell preparation and their purity.

Among the 3259 proteins detected, 612 proteins involved in glucose and lipid metabolism were identified. We performed an unsupervised multivariate analysis focused on these proteins, which clearly demarcated the 4 samples according to their anatomical location (Fig 3A). The first two components of this analysis explained 41% and 20.8% of the dataset variance, respectively (Fig 3A). Statistical analysis of the 612 metabolic proteins differentially expressed in BM-Ad compared to SC-Ad identified 68 up-regulated proteins and 67 down-regulated proteins (Fig 3B). Pathway enrichment analysis with gene analytic software showed clear differences in the expression of proteins involved in several lipid metabolism pathways according to their anatomical locations (Fig 3C). Compared to SC-Ad, BM-Ad showed enrichment in arachidonic acid (AA) metabolism, SL signaling pathway and cholesterol metabolism delineated through cholesterol biosynthesis and statin pathways, while BM-Ad displayed downregulation of glucose and FA metabolism, as well as lipolysis regulation pathways (Fig 3C). Lipoprotein metabolism was also enriched and down-regulated in BM-Ad compared to SC-Ad (Fig 3C). In depth analysis of the proteins differentially expressed in this pathway revealed unexpected specificity for each fat depot.

**Fig 3:**
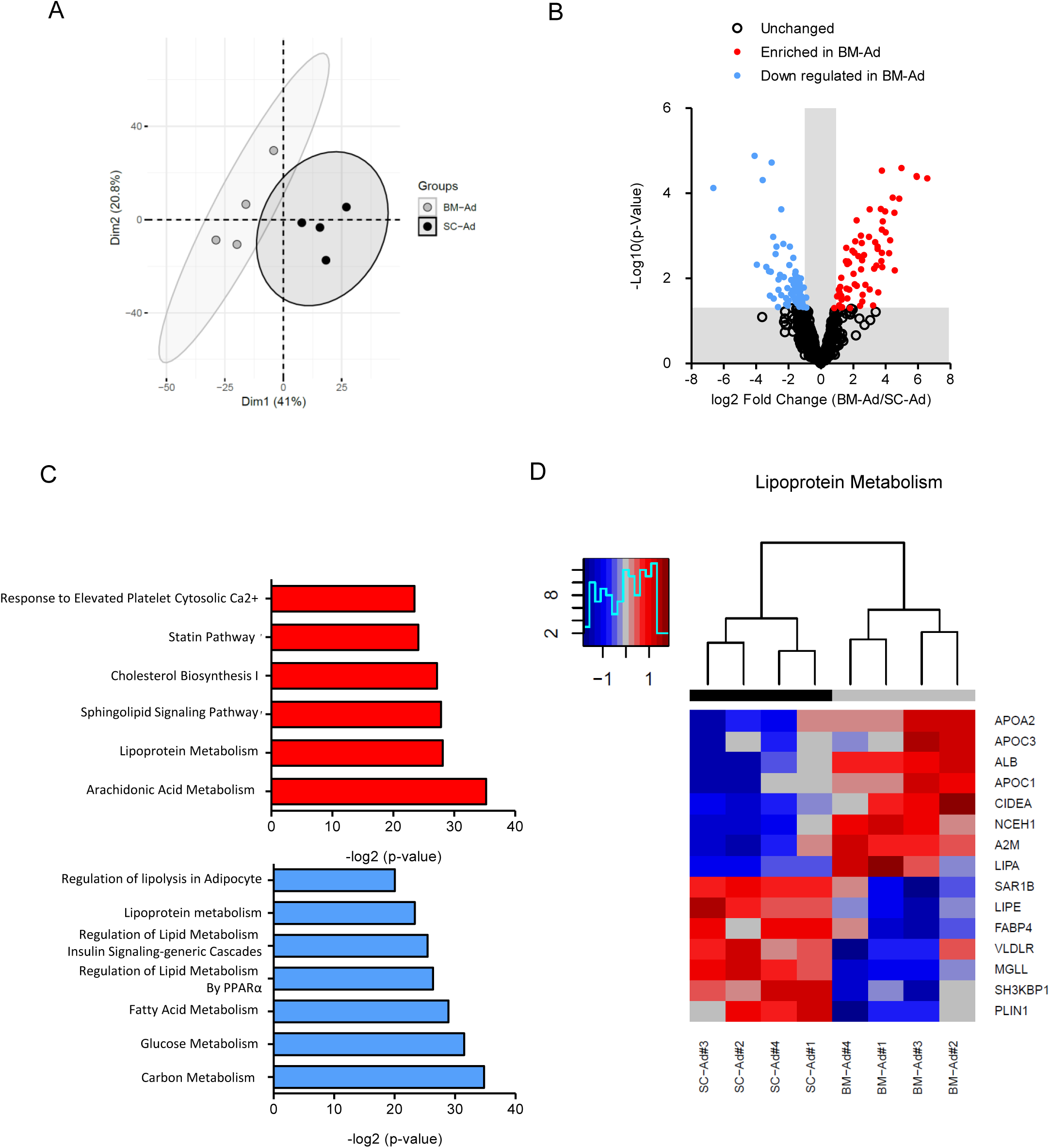
Large-scale proteome analysis highlights differences in lipid metabolism between BM-Ad and SC-Ad. **A.**Principal component analysis of BM-Ad (grey) and SC-Ad (black) based on the relative quantification of the abundance of the proteins involved in lipid and glucose metabolism identified in the LC-MS/MS dataset. The first two components and the percentage of variance for each component are shown. Ellipses show the 95% confidence are interval to strengthen the clustering of the tissues according to their anatomical locations. **B.** Volcano plot of the 612 proteins involved in lipid and glucose metabolism identified and quantified using LC-MS/MS analysis. Sixty seven proteins are significantly (p< 0.05) enriched in BM-Ad (red circles), whereas 68 proteins are significantly (p < 0. 05) down-regulated (blue circle) and 477 are unmodified (open circle) in BM-Ad compared to SC-Ad according to linear model statistical analysis. **C**. Pathway enrichment analysis performed with gene analytics. The top pathways enriched in BM-Ad (red bars) and down-regulated (blue bars) are presented. **D**. Heatmap of the relative abundance of proteins differentially expressed in BM-Ad and SC-Ad belonging to the lipoprotein metabolism pathway. Dendrogram represents hierarchical clustering of the samples. Blue squares represent down regulated proteins and red squares enriched proteins.

We uncovered a clear separation of the samples by their location using hierarchical clustering dendrogram (Fig 3D). In BM-Ad, proteins involved in cholesterol transport (apoliproproteins APO-A2, -C1 and -C4) and hydrolysis, such NCEH1 (Neutral Cholesterol Ester Hydrolysis) and LIPA (Lipase A) (Litvinov et al, 2018), were enriched. This cholesterol-oriented metabolism in BM-Ad is strengthening by the enrichment of several proteins involved in cholesterol biosynthesis and statin pathways (Fig EV3C). On the opposite, proteins related to lipolysis were down-regulated in BM-Ad compared to SC-Ad in lipoprotein metabolism pathway (Fig 3D). Surprisingly, the lipases involved in TG hydrolysis LIPE (gene name of HSL) and MGLL (gene name of MAGL) were decreased (1. 19 ± 0.80, p= 0.016 for LIPE and -2.45 ± 0.35 p =0.00024 for MGLL, see Table EV1), as well as the Fatty Acid Binding Protein 4 (FABP4), one of the most abundant proteins in adipocytes that participate in maintaining adipocyte homeostasis and regulating lipolysis and adipogenesis (Prentice et al, 2019). Finally, BM-Ad and SC-Ad exhibited distinct expression patterns of proteins involved AA metabolism and FA metabolism (Fig EV3D-E). Despite similar morphology and expression pattern of adipokines, our results strongly support that BM-Ad are adipocytes that exhibit a very specific lipid metabolism compared to the “classical” SC-Ad. We uncovered an accumulation of free cholesterol in these cells. This conclusion is supported by unbiased proteomic approaches that indicate a seemingly unidentified cholesterol-oriented metabolism. In contrast to our results, a recent transcriptomic study comparing gene expression of human BM-Ad isolated from the femoral head and SC-Ad show that genes over-represented in human BM-Ad participate in signaling pathways without clear differences in the enzymes involved in lipid metabolism (such as cholesterol metabolism and TG hydrolysis) (Mattiucci et al, 2018). In addition, this report found decreased adiponectin expression, which stand in contrast to our current results and another study that identified BM-Ad as an important source of adiponectin (Cawthorn et al, 2014). We speculate technical issues, such as transcriptomic vs proteomic, and the different sources of BM-AT used may underlie the differences in these findings. We suspect that the specificity of BM-Ad in cholesterol metabolism may reflect their role in supporting BM hematopoiesis (Naveiras et al, 2009; Zhou et al, 2017). Cholesterol is essential constituent of the plasma membrane (Abe & Kobayashi, 2017) and could sustain cell division and plasma membrane fluidity and synthesis of surrounding hematopoietic cells that are under constant renewal. In contrast, the main functions of adipocytes, liberating energy reserve stores as TG under times of energy demand, appears downregulated in these cells. This interesting observation is consistent with the absence of a decrease in BM adiposity under energy deficit conditions (Bathija et al, 1979; Cawthorn et al, 2014; Devlin et al, 2010; Tavassoli, 1974). In particular, we observed a critical down-regulation of MAGL expression, a lipase required for the final hydrolysis of MG produced by HSL activation (Zechner, 2015). As such, MAGL deficiency in mice leads to a concomitant increase in MG levels in AT (Taschler et al, 2011) as observed in BM-Ad. The concomitant decrease of MAGL expression and increase in MG species strongly suggests that MAGL activity may be impaired in BM-Ad compared to SC-Ad.

### Human primary BM-Ad present a defect in lipolytic activity not recapitulated in *in vitro* models

Due to the potential high impact of this newly described regulation in BM-Ad physiology, we further characterized the lipolytic pathway using Western blot analysis of the three major lipases involved in the consecutive hydrolysis of TG. While we observed no differences in ATGL and HSL expression in BM-Ad compared to paired SC-Ad, we found a sharp decrease (about 5-fold) in MAGL (Fig 4A). While we found a slight (1.21-fold) decrease in HSL protein expression in our proteomic studies, this result was not reproduced using Western blot analysis. This discrepancy highlights inter-individual variability. We then functionally assayed lipolytic activity using *ex vivo* approaches on isolated adipocytes. Under basal conditions, we observed reduced glycerol and FFA release in BM-Ad compared to SC-Ad (Fig 4B and 4C). Under isoprenaline stimulation (a β-adrenergic agonist that serves as a strong lipolytic inductor) (Lafontan & Langin, 2009), we found no increase in glycerol (reflecting complete lipolytic reactions) or significant FFA release, but we did find a 3-fold increase in SC-Ad as expected (Lafontan & Langin, 2009). Thus, our data clearly demonstrate that human BM-Ad are devoid of lipolytic activity, which confirms their metabolic specificity.

**Fig 4:**
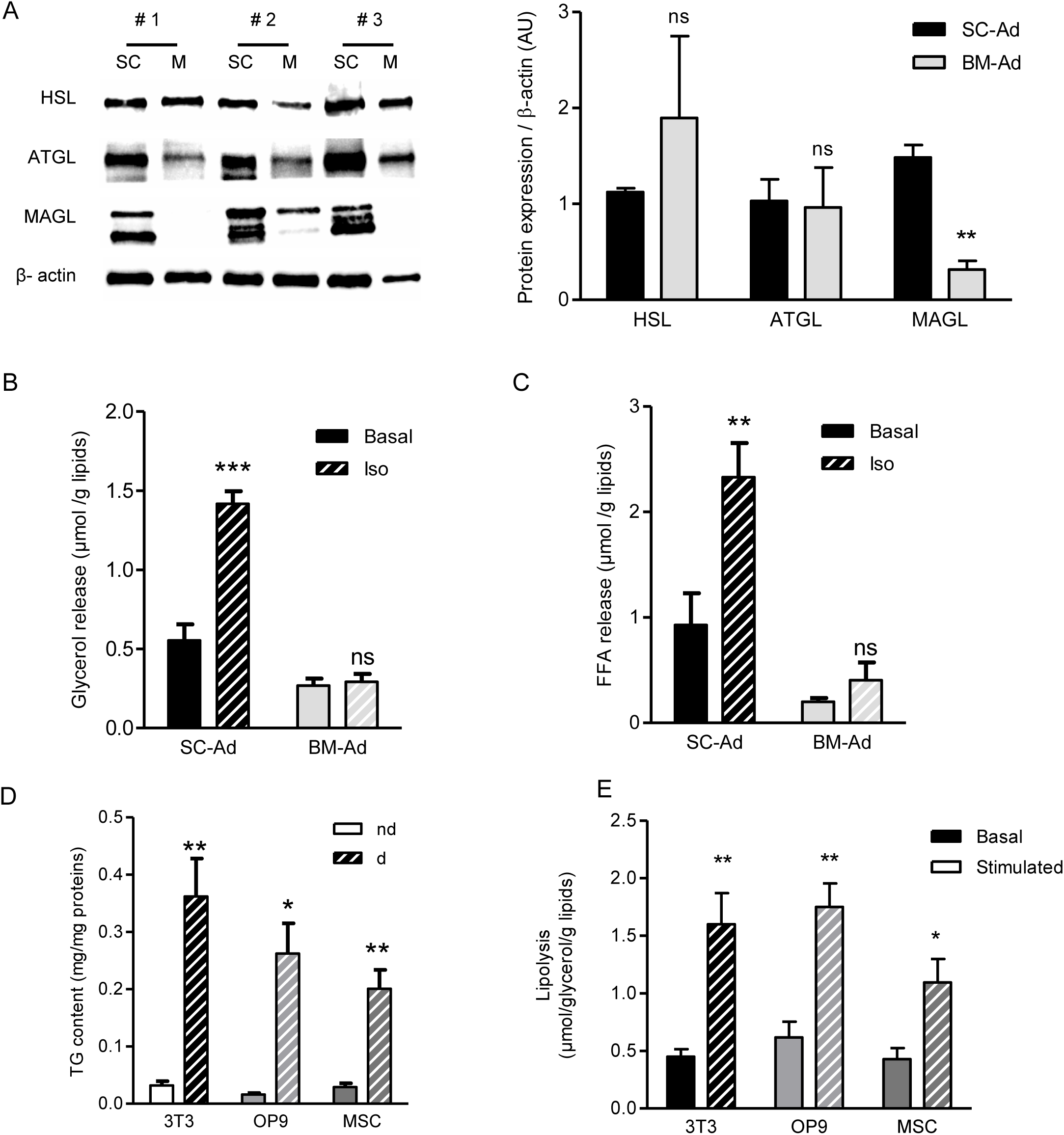
Human native BM-Ad are devoid of lipolytic activity, a metabolic trait not recapitulated by primary BM-MSCs differentiated *in vitro*. **A.**Left panel: Western blot analyses of the three main enzymes involved in lipolysis on paired isolated SC-Ad (SC) and BM-Ad (M) from 3 independent donors. β-actin is shown as loading control. Right panel: relative quantifications of the band intensity normalized to the quantity of β-actin. The histograms represent mean ± SEM. ns=not significant. ** p<0.01 according to paired Student’s t-test. **B.** Glycerol release was measured from isolated SC-Ad and BM-Ad as readout of complete lipolysis under basal condition (plain bar) or after stimulation with isoprenaline (hatched bar). The data are mean of 7 independent experiments and normalized to the quantity of the total lipids content. The histograms represent mean ± SEM, ns, not significant; ***p<0.001 according to two-way ANOVA followed by Bonferroni post-test. **C**. Free fatty acids (FFA) release from isolated SC-Ad and BM-Ad as readout of lipolysis under basal condition (plain bar) or after stimulation with isoprenaline (hatched bar). Data are mean of 7 independent experiments and are normalized to the quantity of the total lipids content. The histograms represent mean ± SEM, ns, not significant; **p<0.01 according to two-way ANOVA followed by Bonferroni post-test. **D**. TG content was measured in cell lysates from 3T3F442A (3T3) and OP9 cell lines or human BM-MSC (MSC) before and after adipogenic differentiation (nd: non-differentiated, d: differentiated). Data are mean of at least 4 independent experiments (4 independent donors were used for human BM-MSC) and were normalized to the quantity of the total protein content. The histograms represent mean ± SEM, *p<0.05; **p<0.01according to two-way ANOVA followed by Bonferroni post-test **E.** Glycerol release from *in vitro* differentiated 3T3F442A (3T3) and OP9 cell lines or human BM-MSC (MSC) as readout of complete lipolysis under basal conditions (plain bar) or after stimulation with isoprenaline (hatched bar). The data are mean of 3 independent experiments (3 independent donors were used for human BM-MSC) and were normalized to the quantity of the total lipids content. The histograms represent mean ± SEM, * p<0.05; **p<0.01 according to two-way ANOVA followed by Bonferroni post-test.

A key finding from our study is the profound down-regulation of MAGL expression, which has never been reported for other AT. Strikingly, the two lipases ATGL and HSL possess several regulators of their activity involving interaction with other proteins as well as phosphorylation state under the control of hormones such as catecholamines (Lafontan & Langin, 2009). However, there is no evidence that cell energy status or hormones can influence MAGL activity, which renders it constitutively active (Lafontan & Langin, 2009). Since BM-Ad down-regulate MAGL expression, this indicates that these cells use the only efficient way to inhibit this specific activity. The slight, but not significant, increase of FFA upon lipolytic stimulation in BM-Ad (Fig 4C) suggests that the lipolysis process is not completely effective in these cells. In mice, a study suggests that rat cBM-Ad (from tail vertebrae) and rBM-Ad (from proximal tibia and femur) are resistant to lipolysis induced by β-adrenergic stimuli. This corresponds at the molecular level to a decrease in active phosphorylation of HSL, whereas the levels of ATGL and HSL remained unchanged compared to SC-Ad (Scheller et al, 2019). Such an additional regulatory process could also occur in human BM-Ad. Interestingly, the MAGL defect in BM-Ad is not compensated by ABHD6 expression, another lipase known to hydrolyze MG in visceral and brown AT but not in SC-AT (Zhao et al, 2016). ABHD6 expression levels remained constant between BM-Ad and SC-Ad in our proteomic study (Table EV1).

We then focused on the metabolic characteristic of BM-Ad we discovered, so we examined the physiological relevance of *in vitro* differentiated adipocytes used as the gold standard model for studying the role of BM-Ad. We differentiated human primary BM-MSCs and murine BM-MSC OP9 cell lines *in vitro* under adipogenic conditions. The murine pre-adipocyte 3T3-F442A served as a control reflecting “classical adipocytes”. In all cells, the differentiation process strongly increased TG content (Fig 4D). *In vitro* differentiated adipocytes from BM-MSCs exhibited similar levels of basal lipolysis compared to differentiated 3T3-F442A (Fig 4E). Isoprenaline stimulation increased glycerol release in all cells studied (Fig 4E). These experiments demonstrated that adipocytes obtained from *in vitro* differentiation of human BM-MSCs do not recapitulate the functional defect in lipolysis observed in BM-Ad isolated from patients. We conclude from these results that *in vitro* differentiated BM-MSCs, considered the gold standard for studying BM-Ad functions, should be interpreted with high caution, since these cells do not recapitulate of key metabolic trait of the BM-Ad phenotype. The differentiation program of BM-Ad may exhibit distinct developmental gene expression patterns and epigenetic signatures that are not induced by the most widely used differentiation protocols that add PPARγ agonists (Lee & Fried, 2014; Ninomiya et al, 2010; Scott et al, 2011). We suggest that such protocols may artificially force BM-Ad progenitors towards a differentiation program into classical white adipocytes.

## Conclusion

Here, we pioneered a methodology to characterize human primary BM-Ad using combined large-scale proteomic and lipodomic approaches. This approach revealed specific markers in a phenotype that refines and identifies BM-Ad. These cells morphologically resemble classical SC-Ad; however, we unraveled the specific lipid metabolism of BM-Ad, including the presence of a cholesterol-orientated metabolism that requires further investigation. We demonstrated altered lipolytic function in human primary BM-Ad due to a profound down-regulation of MAGL expression. This result underlies the differences in metabolic fitness upon caloric restriction between BM-AT and SC-AT. This specific phenotype is a previously unidentified feature of adipose depots that could explain why BM-AT behaves like a preserved lipid source, except during periods of extremely severe nutrient deprivation (Abella et al, 2002; Cawthorn et al, 2016). The specific function of this preservation, whether overall metabolic fitness or local interaction with proximal cells (such as hematopoietic cells), remains unknown. Adipocyte diversity continues to increase, so distinguishing markers and delineating specific phenotypes of these adipocyte subtypes gain importance. In addition to white and brown adipocytes, recent studies have identified beige adipocytes, an inducible form of thermogenic adipocytes (Zwick et al, 2018) and pink adipocytes in mouse mammary fat pad during pregnancy and lactation (Giordano et al, 2014). Given their specificity for lipid metabolism regardless of their morphological similarity to white adipocytes, we propose to define BM-Ad as a distinct type of adipocytes named “yellow adipocytes”.

## Materials and Methods

### Human subcutaneous and bone marrow tissue samples

Paired subcutaneous (SC-AT) and bone marrow adipose tissue (BM-AT) were harvested from patients undergoing hip surgery in the Orthopedic Surgery and Traumatology Department of the Hospital Pierre Paul Riquet (Toulouse, France). All patients gave their informed consent and the samples were obtained according to national ethic committee rules (authorization° DC-215-2342). Briefly, during total hip replacement surgeries, after skin incision, maximus gluteus muscle and external rotators dissection, an osteotomy of the femoral neck was performed which allowed access to the intramedullary canal. While broaching the canal, BM-AT was aspirated cautiously with a soft cannula in the femoral proximal metaphysis and diaphysis before prosthesis placement. All procedures were performed using the same posterior approach. SC-AT were harvested using surgical blade at the incision site in the gluteal region. The samples were immediately placed in 37°C pre-warmed KRBHA (Krebs Ringer BSA HEPES Albumin buffer) corresponding to Krebs Ringer buffer (Sigma-Aldrich) supplemented with 100mM Hepes (Sigma-Aldrich) and 0.5% free fatty acid (FFA-free) bovine serum albumin (BSA) (Sigma-Aldrich) and rapidly carried out to the laboratory (within 1h) where they were processed. The BM-AT that share the same macroscopic aspects compared to SC-AT was dissected from area rich in hematopoietic cells (red marrow). For all the experiments performed in our study, 24 independent samples were used and obtained from 14 men and 10 women (mean age: 66.7 ± 13.9 years and mean body mass index (BMI): 26.8 ± 3.4 kg/m^2^).

### Adipocyte isolation

SC-AT and BM-AT were rinsed several times in KRBHA prior to collagenase digestion (250 UI / mL diluted in PBS calcium and magnesium free supplemented with 2% FFA-free BSA (all products were obtained Sigma-Aldrich). After 30 min digestion at 37°C under constant shaking, samples were filtered with 100 and 200µm cell strainers (for BM-AT and SC-AT respectively) to remove cellular debris, undigested fragments and bone trabeculaes. The cell suspension was then gently centrifuged for 5 min at 200g at room temperature (RT). The floating adipocytes were then collected and rinsed with KRBHA several times to obtain a pure adipocyte cell suspension.

### SC-AT and BM-AT confocal microscopy

Pieces of 0.5 cm^2^ of whole SC-AT and BM-AT were fixed with a 4% paraformaldehyde solution (PFA, Electron Microscopy Sciences (EMS)) overnight. Fixed tissues were blocked and permeabilized in calcium and magnesium free PBS supplemented with 3% BSA and 0.2% Triton X100 (both obtained from Sigma Aldrich) for 1 h at RT. Tissues were then incubated overnight with a mouse anti PLIN1 serum (Acris Biosystem; 1:10 in calcium and magnesium free-PBS, 3% BSA, 0.2% Triton X-100). The following day the tissues were rinsed 5 times in PBS 0.05% Tween-20 and incubated for 2 h with a secondary antiboby coupled with CF488 dye (Biotum) for PLN1 staining, rhodamine coupled phalloidin (Thermofisher) for filamentous actin staining and TOPRO3^®^ (Thermofisher) for nuclei staining. Z-stack images were acquired using LSM 710 confocal microscope and a 10× or 40× objective (Zeiss). Maximum intensity projection was made using Image J software and orthogonal views using Imaris software (v9.2; Bitplane).

### Transmission electron microscopy

SC-AT and BM-AT were fixed with 2,5% glutaraldehyde and 2% PFA (EMS, Hatfield, PA, USA) in Cacodylate buffer (0.1M, pH 7.2) overnight at 4 °C and post-fixed at 4°C with 1% OsO4 and 1.5% K3Fe(CN)6 in the same buffer. Samples were treated for 1 h with 1% aqueous uranyl acetate and were then dehydrated in a graded ethanol series and embedded in EMBed-812 resin (EMS). After 48 h of polymerization at 60 °C, ultrathin sections (80 nm thick) were mounted on 75 mesh formvar-carbon coated copper grids. Sections were stained with 2% uranyl acetate (EMS) and 3% Reynolds lead citrate (Chromalys). Grids were examined with a TEM (Jeol JEM-1400, JEOL Inc) at 80 kV. Images were acquired using a digital camera (Gatan Orius, Gatan Inc, Pleasanton, CA, USA).

### Confocal microscopy on isolated adipocytes

BM-Ad and SC-Ad were isolated as described above. Immediately after isolation, primary adipocytes were embedded in a fibrin gel to maintain cellular integrity. Briefly, 30µl of isolated adipocytes were gently mixed with 30µl of a fibrinogen solution (18 µg/mL in 0.9% NaCl buffer, Sigma-Aldrich) and 30µl of thrombin (3 units in 30µl of CaCl_2_ solution, Sigma-Aldrich). Gel polymerization occurs rapidly at 37°C. The gels containing the primary adipocytes were fixed in 4% PFA for 1h and incubated with 10 ng/mL of Bodipy^®^ 493/503, rhodamine coupled phalloidin and TOPRO3 (all products were obtained from Thermofischer). Samples were examined using LSM 710 confocal microscope and a 40X objective (Zeiss). Maximum intensity projection was performed using Image J software.

### Lipidomic analysis

For the lipidomic and proteomic studies, 4 samples were used (2 men and 2 women, mean age: 67± 7. 4 years; mean BMI: 26.5 ± 3.1 kg/m^2^). After 3 washes with PBS, isolated adipocytes (400 µl) were mixed 1.5 mL methanol in glass tubes. Sample mixture was then incubated with 5 mL of methyl tert-butyl ether (MTBE ;Sigma-Aldrich) for 1h at RT under gently shaking. After adding 1.2 mL of water, samples were centrifuged for 10 min at 1000 g. Upper phase (containing lipids) was transferred in a new glass tube. Lower phase was re-extracted with 2 volume parts of MTBE: methanol: water (10: 3: 2.5) and samples were centrifuged for 10 min at 1000 g and used for proteomic analyses (see below). Upper phase was collected, combined with the one collected after the first extraction and kept at -80°C for lipidomic analysis. One ml of lipid phase was evaporated under a nitrogen stream. Dried samples were sent to the Harvard mass spectrometry core and were analyzed by their untargeted lipidomics profiling platform. Lipids were resuspended with 100µl of 1:1 LC/MS grade isopropanol: acetonitrile methanol and 5 µl were injected onto the LC-MS/MS. Data acquisition was performed as previously described (Breitkopf et al, 2017). Briefly, each peaks area was calculated in both positive and negative ionization mode. The peaks allowing to structurally resolving the same lipid species were sum, if obtained from the same ionization. Only the lipid species detected at least in 3 samples from the same location were considered as robustly detected and used for the analysis. Missing values were imputed as the first percentile of the entire dataset. Then, values were log2 transformed and normalized with the function NormalizeBetweenArrays from Bioconductor package to perform the principal component analysis with R software (v3.5) and FactomineR package. The heatmaps and associated hierarchical clustering build on K-means methods were resolved with R software and ggplot2 package after centering the data around zero. The lipid species belonging to the same classes were sum to measure their relative abundance and the log2 fold change of signal intensities for each class was calculated to compare the lipid classes between adipocyte locations. Violin plot was drawn with vioplot function in R.

### Cholesterol content quantification

Cholesterol content within isolated adipocytes was measured using cholesterol assay kit (obtained from Abcam-ab65390) according to manufacturer recommendations. Briefly, lipids were extracted from isolated adipocytes using MTBE as described above. Free cholesterol and total cholesterol was sequentially quantified using colorimetric method. Optical density was determined at 570 nm with µ-quant spectrophotometer (BioTek Instruments).

### Proteomic analysis

After 3 washes with PBS, proteins from isolated adipocytes (400µl) were purified with 5 mL of MTBE as described in lipidomic analysis section. Lower phase was centrifuged for 10 min at 5000 g at RT and pellet (containing proteins) was washed 2 times with PBS to remove solvents. Pellets were then resuspended with PBS 2% SDS, sonicated for 20 seconds and protein concentration was determined with the commercial kit (DC™ Protein Assay; Bio-Rad). Fifteen µg of proteins were reduced with modified Laemmli buffer (40 mM Tris pH 6.8, 2% SDS, 10% glycerol, 25mM DTT and 0.01% bromophenol blue) for 15 min at 65°C and alkylated by addition of 90mM iodoacetamide for 30 min at RT in the dark. Protein samples were loaded on a 1D SDS-PAGE gel (0.15 × 8 cm) and the electrophoretic migration was stopped as soon as the proteins entered the separating gel, in order to isolate all proteins in a single gel band (stained with Coomasie blue). The corresponding gel slice was excised and washed with 100 mM ammonium bicarbonate buffer. Proteins were in-gel digested using 0.6 µg of modified sequencing grade trypsin (Promega) in 50 mM ammonium bicarbonate overnight at 37°C. The resulting peptides were extracted in 50 mM ammonium bicarbonate followed by 10% formic acid/acetonitrile (1/1 v/v). The peptidic fractions were dried under speed-vacuum and resuspended with 5% acetonitrile, 0.05% trifluoroacetic acid (TFA) for MS analysis.

Peptides were analyzed by nanoLC-MS/MS using an UltiMate 3000 RSLCnano system coupled to a Q-Exactive Plus mass spectrometer (Thermo Fisher Scientific, Bremen, Germany). Five µL of each sample were loaded on a C-18 precolumn (300 µm ID × 5 mm, Thermo Fisher) in a solvent made of 5% acetonitrile and 0.05% TFA and at a flow rate of 20 µL/min. After 5 min of desalting, the precolumn was switched online with the analytical C-18 column (75 µm ID × 15 cm, Reprosil C18) equilibrated in 95% solvent A (5% acetonitrile, 0.2% formic acid) and 5% solvent B (80% acetonitrile, 0.2% formic acid). Peptides were eluted using a 5 to 50% gradient of solvent B over 300 min at a flow rate of 300 nL/min. The Q-Exactive Plus was operated in a data-dependent acquisition mode with the XCalibur software. MS survey scans were acquired in the Orbitrap on the 350-1500 m/z range with the resolution set to a value of 70000. The 10 most intense ions per survey scan were selected for HCD fragmentation and the resulting fragments were analyzed in the Orbitrap with the resolution set to a value of 17500. Dynamic exclusion was employed within 30 seconds to prevent repetitive selection of the same peptide. Duplicate technical LC-MS measurements were performed for each sample.

Raw mass spectrometry files were processed with the MaxQuant software (version 1.6.3.4) for database search with the Andromeda search engine and quantitative analysis. Data were searched against human entries of the Swissprot protein database (UniProtKB/Swiss-Prot Knowledgebase release 2018_02). Carbamidomethylation of cysteines was set as a fixed modification whereas oxidation of methionine and protein N-terminal acetylation were set as variable modifications. Specificity of trypsin digestion was set for cleavage after K or R, and two missed trypsin cleavage sites were allowed. The precursor mass tolerance was set to 20 ppm for the first search and 4.5ppm for the main Andromeda database search. The mass tolerances MS/MS mode was set to 0.5 Da. Minimum peptide length was set to seven amino acids, and minimum number of unique peptides was set to one. Andromeda results were validated by the target-decoy approach using a reverse database at both a peptide and a protein FDR of 1%. For label-free relative quantification of the samples, the “match between runs” option of MaxQuant was enabled with a time window of 0.7min, to allow cross-assignment of MS features detected in the different runs.

The “LFQ” metric from the MaxQuant “protein group.txt” output was used to quantify proteins. Missing protein intensity values were replaced by a constant noise value determined independently for each sample as the lowest value of the total protein population. Only proteins identified in at least three samples in the same location (i.e. SC-Ad or BM-Ad) were considered as robustly detected and were used for statistical and bioinformatic analyses. Protein involved in lipid and glucose metabolism were selected using gene analytics software based on their involvement into the following pathways: Regulation of lipid metabolism; Insulin signaling-generic cascades; Lipoprotein metabolism; Adipogenesis; Regulation of lipid metabolism by Peroxisome proliferator-activated receptor alpha; Glucose / Energy Metabolism; Peroxisomal lipid metabolism; Calcium (Ca), cyclic adenosine monophosphate (cAMP) and Lipid Signaling; Nuclear Receptors in Lipid Metabolism and Toxicity; SREBF (Sterol Regulatory Element Binding Protein Gene) and miR33 in cholesterol and lipid homeostasis; Acyl chain remodeling of Phospho Ethanolamine (PE) ; Cholesterol and Sphingolipids transport / Distribution to the intracellular membrane compartments; Synthesis of substrates in N-glycan biosynthesis; Synthesis of Phosphatidyl Choline (PC); Metabolism of steroid hormones; Glycerophospholipid biosynthesis; Glucose metabolism; Fat digestion and absorption; Regulation of cholesterol biosynthesis by SREBP (Sterol Regulatory Element Binding Protein); cholesterol biosynthesis III (via desmosterol); Cholesterol and Sphingolipids transport / Transport from Golgi and ER to the apical membrane; Aldosterone synthesis and secretion; Citrate cycle (Tricarboxylic Acid (TCA) cycle); Adipocytokine signaling pathway; Sphingolipid metabolism; Fatty acid metabolism; Pyruvate metabolism; Arachidonic acid metabolism; Linoleic acid metabolism; Ceramide Pathway; Sphingolipid signaling pathway; sphingomyelin metabolism/ceramide salvage; Pentose phosphate pathway; Regulation of lipolysis in adipocytes; Mitochondrial Long Chain-Fatty Acid, Beta-Oxidation SuperPath; Fatty acid biosynthesis. Among the 1948 proteins retrieved by the database, we robustly identified 612 proteins. The label free quantification (LFQ) intensity for each identified protein was log2 transformed and used to perform the principal component analysis with R software (v3.5) and FactomineR package. The statistical analysis of differentially expressed proteins was performed with LIMMA package from Bioconductor using linear model followed by borrowing strength across protein with empirical bays methods with a design matrix build on two groups (BM-Ad and SC-Ad) (see Supplementary Table 1). The protein expression was considered significantly different if the p-value was lower than 0.05. Pathway enrichment analysis was performed with gene analytics software. The official gene symbol of the proteins significantly enriched or down-regulated was used as entry to determine the pathways enriched or downregulated respectively. The heatmaps and associated hierarchical clustering build on K-means methods were resolved with R and ggplot2 package.

### *In vitro* adipogenesis

The pre-adipocyte 3T3 F442A obtained from ECACC (00070654) were grown and differentiated into adipocyte as previously described (Meulle et al, 2008). OP9 cell were obtained from ATCC (ATCC CRL-2749). OP9 cells were seeded at 1×10^5^ cell/well in 6-well plates for 2 days in MEM alpha supplemented with 20% fetal calf serum (FCS), 125 mg/mL streptomycin, 125 UI/mL penicillin. At 80% of confluence, media was replaced with similar media supplemented with 15% knock-out serum (invitrogen10828-028) for 5 days to induce adipogenic differentiation (Wolins et al, 2006). Human BM-MSC were isolated from bone marrow (obtained by sternal puncture) of healthy patients as previously described (Corre et al, 2007). BM-MSC (passage 2) were seeded at 3×10^5^ cell/well in 6-well plates for 2 days in MEM alpha supplemented with 10% fetal calf serum (FCS), 125mg/mL streptomycin, 125UI/mL penicillin. At 80% of confluence, media was replaced with StemMACS™ AdipoDiff Media (Miltenyi 130-091-677) supplemented with 125mg/mL streptomycin, 125UI/mL penicillin to induce adipogenic differentiation for 28 days. Media was changed every 2 to 3 days and cells were grown in a humid atmosphere with 5% CO2 at 37°C. TG content of cells before and at the end of adipogenic differentiation was performed as previously described (Dirat et al, 2011) using commercial kit (Sigma-F6428).

### Western blot

Isolated adipocytes were washed 3 times with PBS and proteins were separated from lipids using MTBE extraction described above (proteomic analysis section). Five µg of proteins were reduced with modified Laemmli buffer for 15 minutes at 65°C, loaded on 4-10% gradient SDS-PAGE gel (Biorad) and transferred to nitrocellulose membrane. Membranes were blocked with 5% skimmed milk in TBS (20mM Tris, 150mM NaCl) and incubated with appropriate primary antibodies (rabbit polyclonal antibody (pAb) anti ATGL, (1/:1000, ref: 2138, Cell Signaling Technology); rabbit pAb anti HSL (1:1000, ref: 4107, Cell Signaling Technology); rabbit pAb anti MAGL (1:1000, ref: sc134749, Santa Cruz Biotechnology); mouse monoclonal anti β-Actin (1:5000, clone: AC-15, Sigma Aldrich). The membranes were washed with TBS complemented with 0.1% Tween-20 and incubated with HRP conjugated secondary antibodies (1:5000, Santacruz Biotechnology). The immunoreactive protein bands were revealed by ECL prime Western blotting detection reagent (Ammersham™) and detected using ChemiDoc™ Imaging System (Biorad). Densitometry quantification was performed using image lab software (v5.2.1; Biorad). Signal intensity was normalized to β-Actin.

### Lipolysis assay

Isolated adipocytes (50μl) were incubated with 450 μL KRBHA with or without isoprenaline 10^-6^ mol. L^-1^ (Sigma Aldrich) to evaluate stimulated and basal lipolysis respectively. After 2 h incubation at 37°C under gentle shaking, 200 μL of incubation media was removed and kept to measure glycerol and FA release using commercial kits (Sigma-F6428 and Wako diagnostic NEFA-HR, respectively). Results were normalized to total lipid content quantified after Dole extraction. Briefly, isolated adipocytes were lysed by the addition of Dole’s Reagent (40:10:1 isopropanol : heptane : H_2_SO_4_ 1N). Upper phase containing lipids was extracted again with heptane, evaporated under a nitrogen stream and dried lipids were weighted. For lipolysis experiment on adipocyte-differentiated cell lines (3T3 F442A and OP9) and human BM-MSC, cells were incubated for 3 hours and results were normalized to TG content. At the end of the incubation, cells were washed with PBS and resuspended in buffer containing 10mM Tris HCL pH 7.5 and 1mM EDTA to quantify TG.

### Statistical analyses

Statistical analyses were performed using Prism v4 (GraphPad Software). Comparison between two groups was performed using paired Student’s t-test and multiple comparisons was performed by two-way ANOVA follow by Bonferroni post-test for n independent experiments. P-value was considered significant if lower than 0.05.

## Acknowledgements

This work benefited from the assistance of Stephanie Balor and Vanessa Soldan from the Multiscale Electron Imaging platform (METi) of the Centre de Biologie Intégrative (Toulouse, France). Lipidomic analysis was performed at the Mass Spectrometry Facility of the Beth Israel Deaconess Medical Center (Boston, USA). This work by supported by the “Fondation de France (contract N° 171352) for running costs and a two-year post-doctoral fellowship for Camille Attané. David Estève received a post-doctoral fellowship from the Fondation pour La Recherche Médicale (SPF201809007124). This work also benefited from the Toulouse Réseau Imagerie (TRI)-RIO Optical Imaging Platform at the Institute of Pharmacology and Structural Biology (Genotoul, Toulouse, France) supported by grants from the Région Midi-Pyrénées (contrat de projets état-région), the Grand Toulouse community, the Association pour la Recherche sur le Cancer (Equipement 8505), the CNRS and the European Union through the Fonds Européen de Développement Régional program. We thank Life Science Editors for editorial assistance.

## Author contributions

NR set up the conditions for harvesting BM-AT and SC-AT in close collaboration with CA and DE and supervised the samples collection. CA, DE, MM handled the AT samples and isolated adipocytes. DE performed the transmission electron microscopy (with the help of the METi platform) and the immunofluorescence experiments as well as image data analysis. CA performed sample preparation for proteomic and lipidomic studies, Western blot, cell culture (with the help of MM) and the lipolysis experiments. JC performed the isolation of human BM-MSC. KC performed the proteomics studies under the supervision of OS. DE and CA conducted analysis of lipidomic (with the help of PV) and proteomic data under the supervision of JI. CA, DE, PV, OS and CM analyzed the data. CA, DE and CM conceived the idea for this project and wrote the manuscript with significant inputs from all authors. CM supervised the study.

## Conflict of interest

The authors declare they have no conflict of interest

## Expanded View Figure legends

**Fig EV1:**
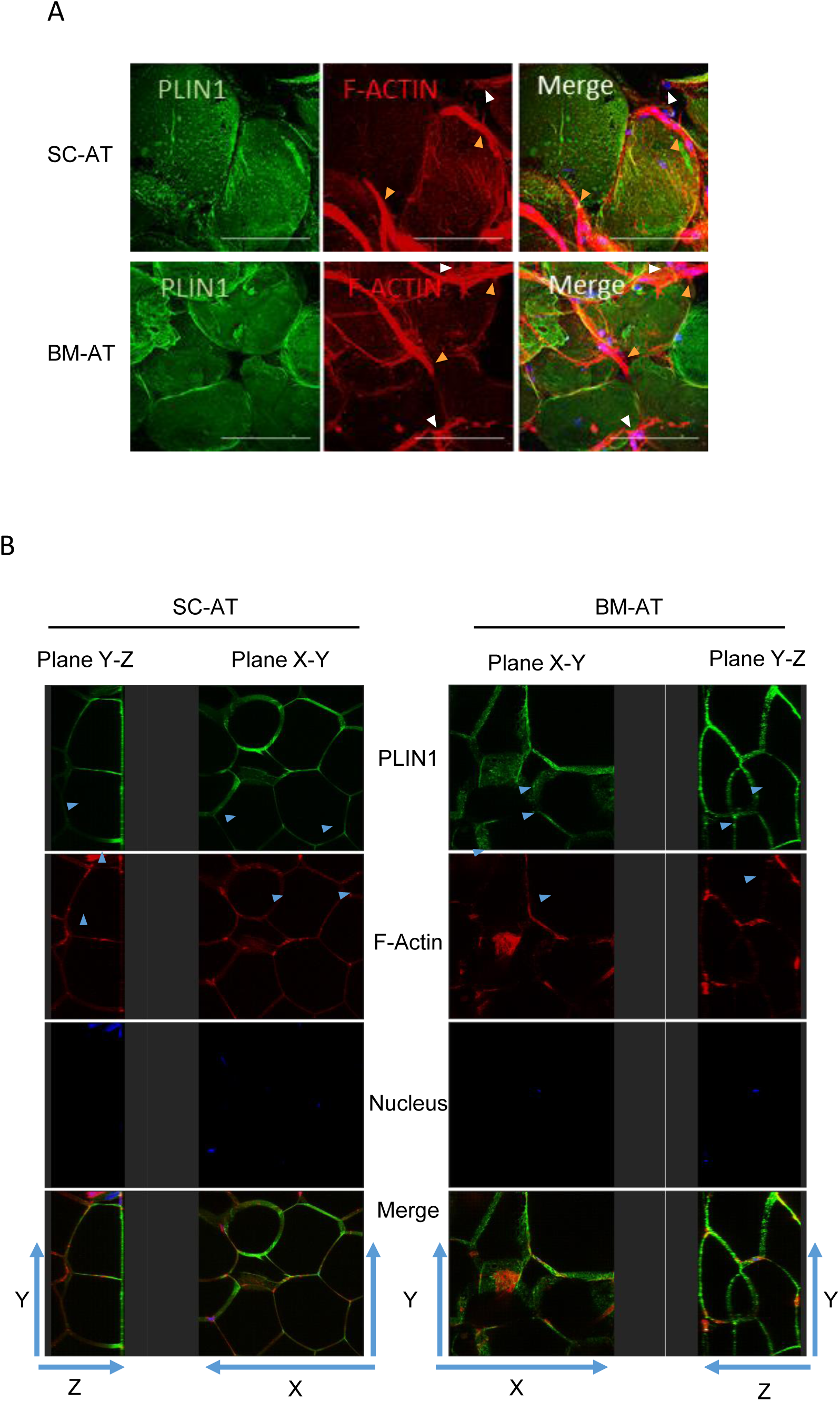
SC-Ad and BM-Ad exhibit similar morphology. **A.** Whole mount SC-AT and BM-AT were stained with an antibody directed against perilipin 1 (PLIN1, green), phalloidin (F-Actin, red) and TOPRO-3 (nucleus, blue). Z stack images were taken using confocal microscope with 40X objective (n=3). Representative maximum intensity projection is shown. Orange arrowheads show vessels and white arrowheads show stromal cells. Scale bar=50µm. **B.** Representative XY and YZ focal planes are shown to highlight the cohesive organization of the SC-and BM-AT and the thin cytoplasm of adipocytes (blue arrowheads).

**Fig EV2:**
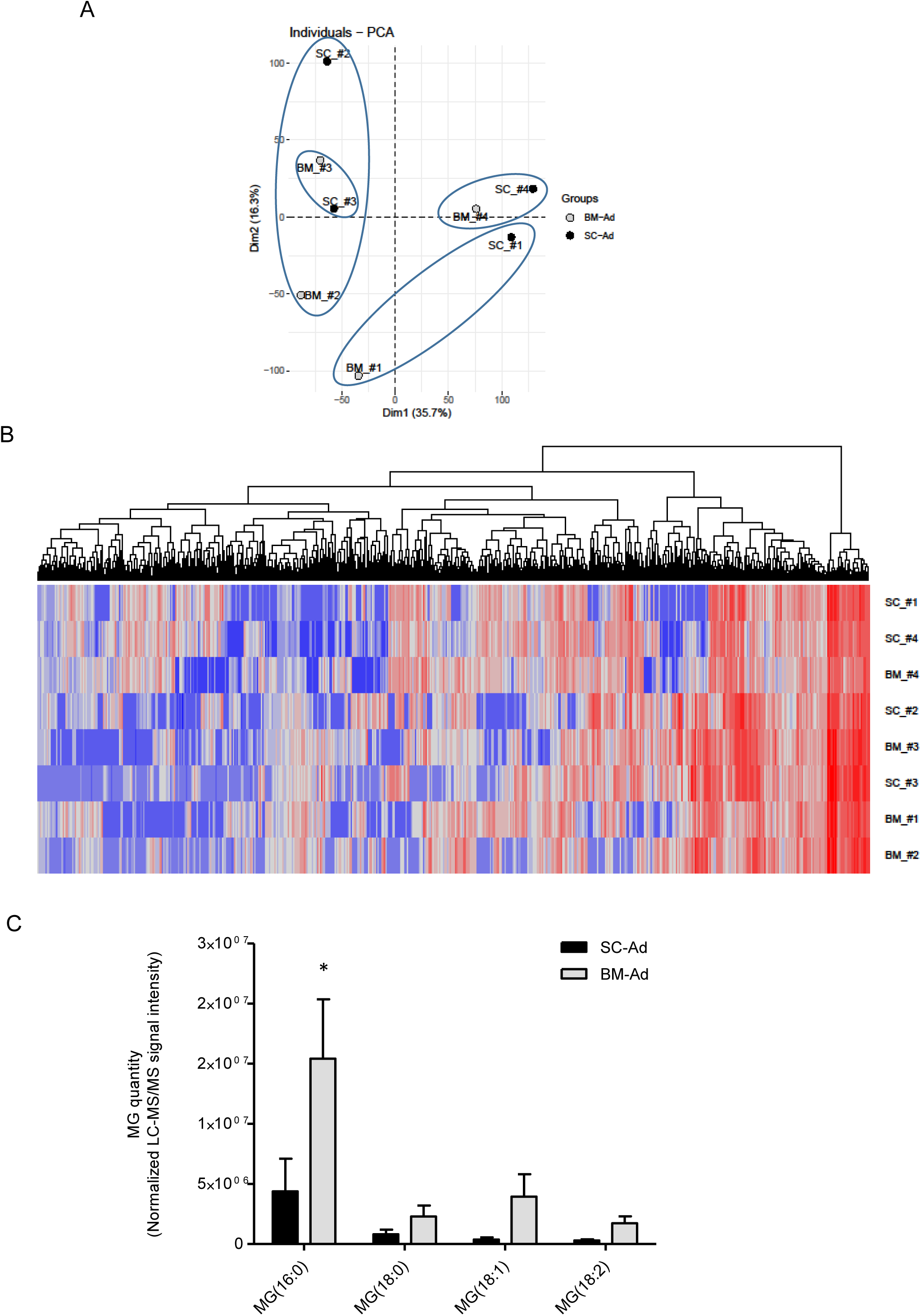
Unsupervised lipidomic analyses reveal that the variance between samples mainly occur through inter-individual variability despite increased MG species in BM-Ad. **A.**Principal component analysis of BM-Ad (grey) and SC-Ad (black) based on the relative quantification of the lipid species identified in LC-MS/MS. The first two components and the percentage of variance for each component are shown. **B.** Heatmap of the relative abundance of lipid species quantified in BM-Ad and SC-Ad. Dendrogram represents hierarchical clustering of samples. Blue squares represent down-regulated lipid species and red squares enriched lipid species. **C**. Relative quantification of the main MG species by LC-MS/MS in paired isolated SC-Ad and BM-Ad (n= 4). Histograms represent mean ± SEM, * p<0.05 according to two way ANOVA followed by Bonferroni’s post-test.

**Fig EV3.**
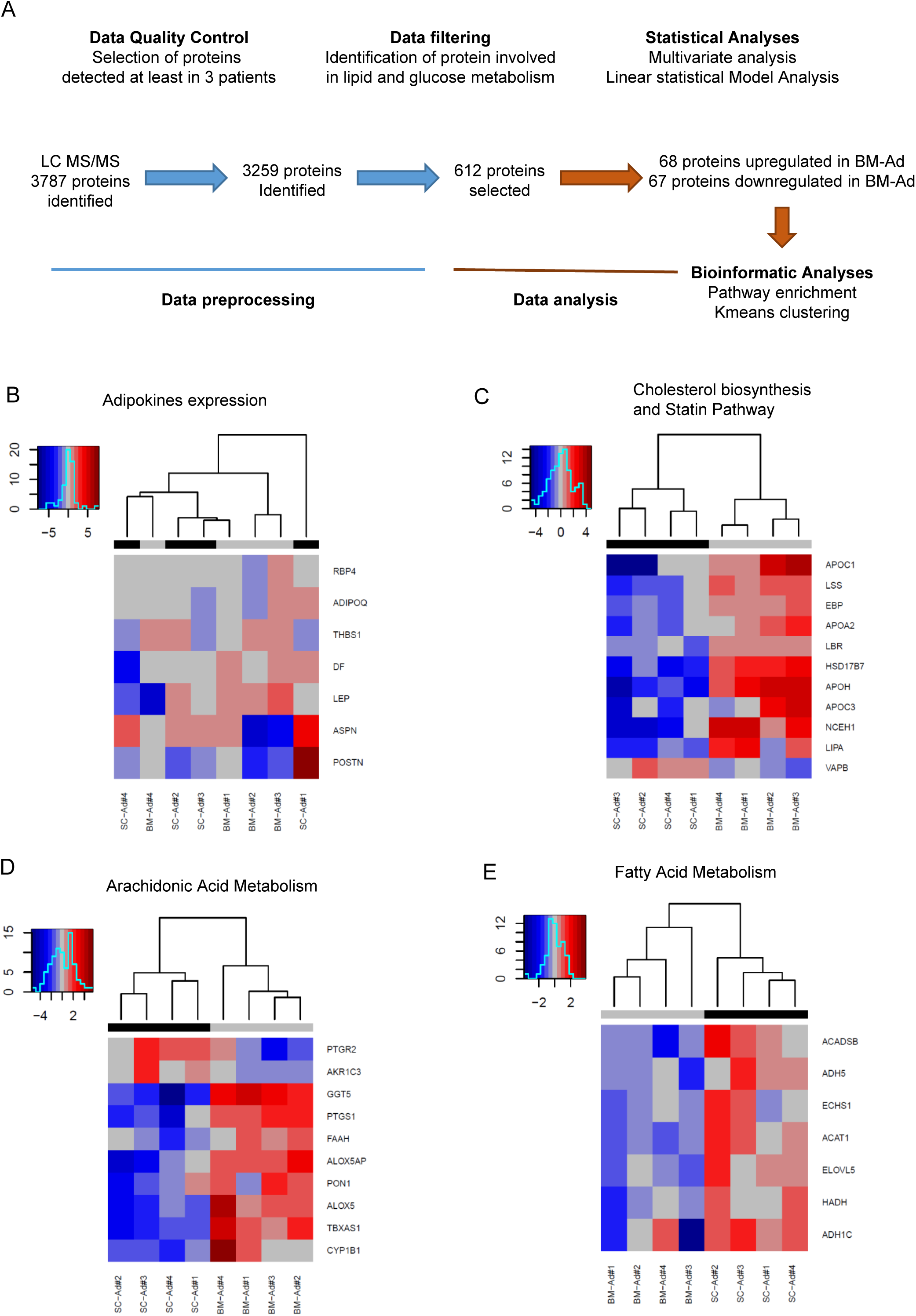
Large-scale analysis of the proteome reveals differences in lipid metabolism, but not adipokines, between BM-Ad and SC-Ad. **A.** Scheme of the proteomic dataset analysis workflow. Extracted proteins from paired isolated SC-and BM-Ad were analyzed by LC-MS/MS. Among the 3787 proteins identified, 3259 were robustly identified in at least 3 of 4 donors. We selected a set of 612 proteins involved in lipid and glucose metabolism using gene analytics software to perform the statistical and bioinformatics analyses. In this dataset, unsupervised multivariate analysis was performed and differentially expressed proteins were identified using a linear statistical model (LIMMA) allowing to identify 67 proteins enriched and 68 proteins downregulated in BM-Ad compared to SC-Ad. Pathway enrichment analyses were performed using gene analytics software that concatenates several databases to identify specific lipid pathways enriched and down regulated in BM-Ad. Hierarchical clustering analyses were then performed on these specific lipid pathways. **B.** Heatmap of the relative abundance of adipokines expressed in BM-Ad and SC-Ad. Dendrogram represents hierarchical clustering of the samples. Blue squares represents down regulated proteins and red squares enriched proteins. **C**. Heatmaps of the relative abundance of proteins differentially expressed in BM-Ad and SC-Ad belonging to the Cholesterol Biosynthesis I and Statin pathway. Dendrogram represents hierarchical clustering of the samples. Blue squares represent down regulated proteins and red squares enriched proteins. **D**. Heatmap of the relative abundance of proteins differentially expressed in BM-Ad and SC-Ad belonging to the arachidonic acid metabolism pathway. Dendrogram represents hierarchical clustering of the samples. Blue squares represents down regulated proteins and red squares enriched proteins. **E.** Heatmap of the relative abundance of proteins differentially expressed in BM-Ad and SC-Ad belonging to the FA metabolism pathway. Dendrogram represents hierarchical clustering of the samples. Blue squares represent down regulated proteins. Red squares enriched proteins.

**Supplementary Table 1:**
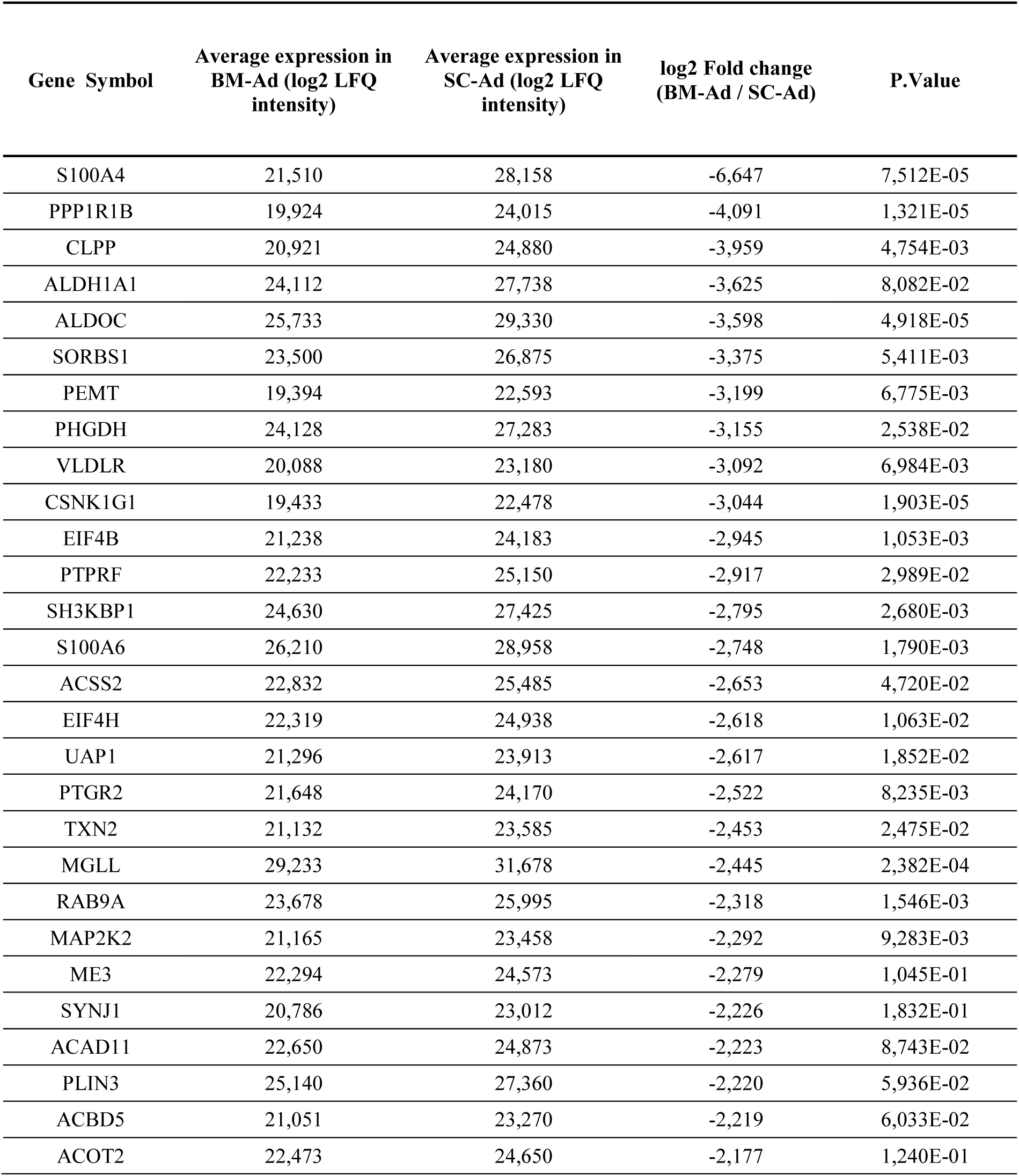
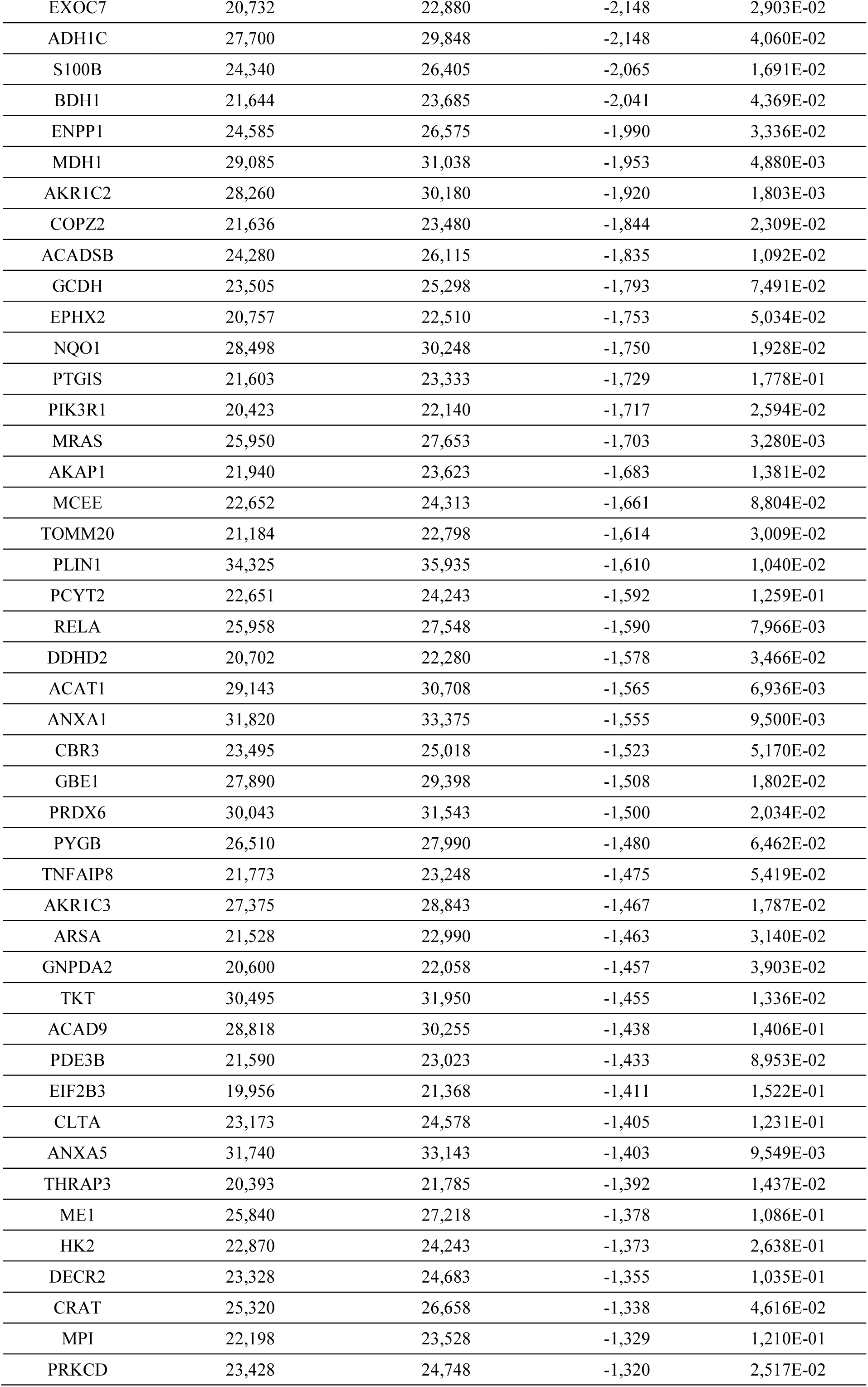
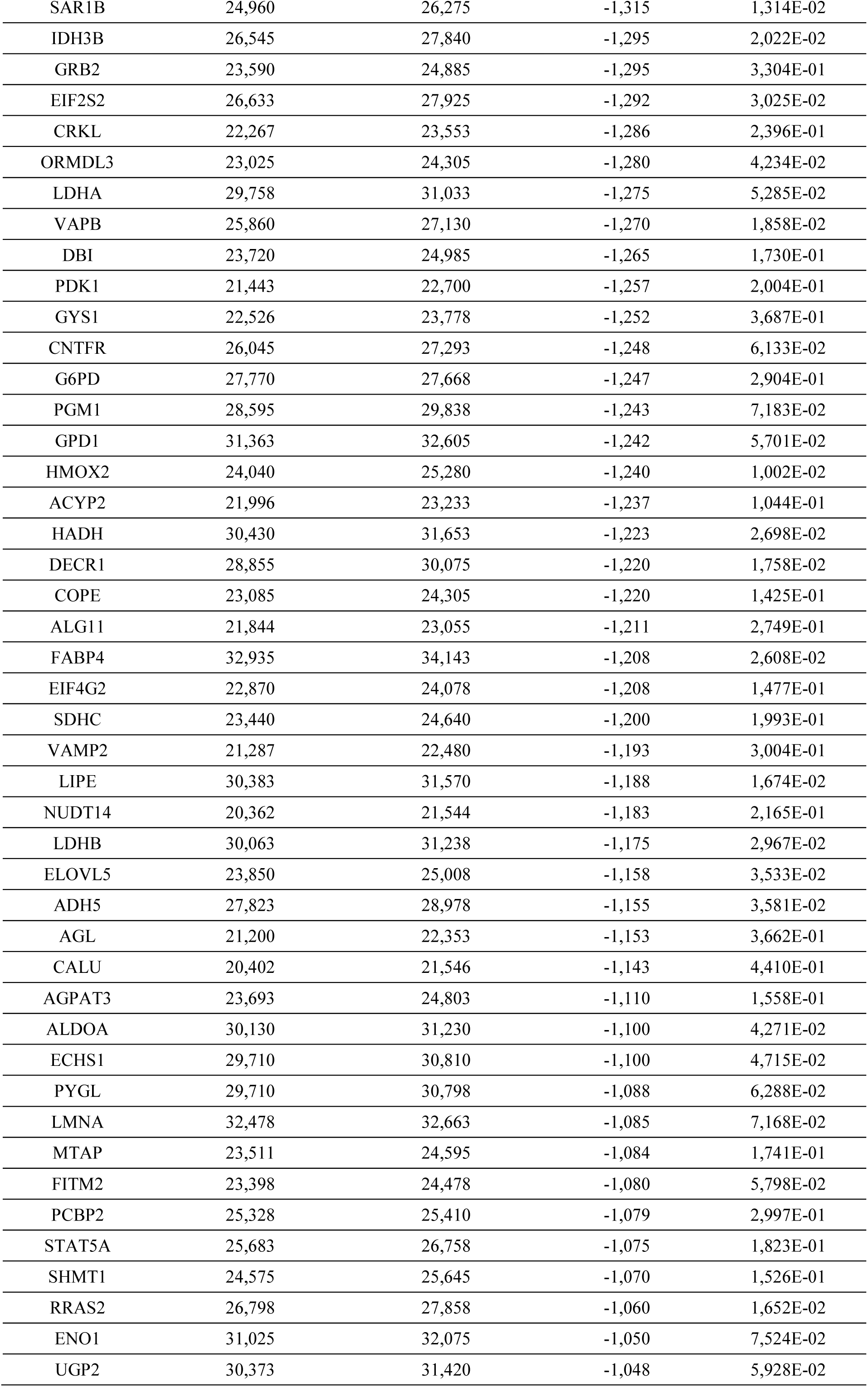
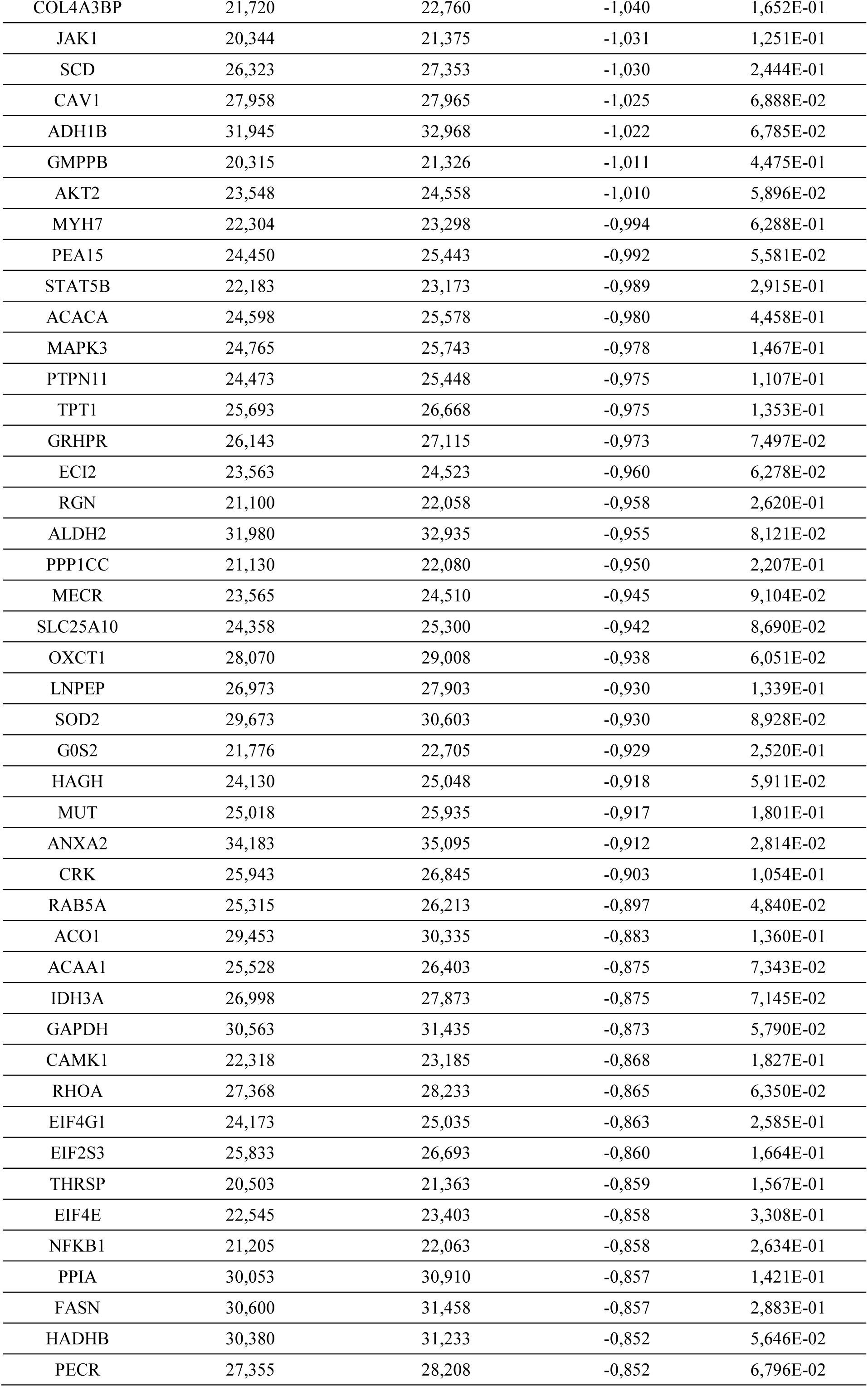
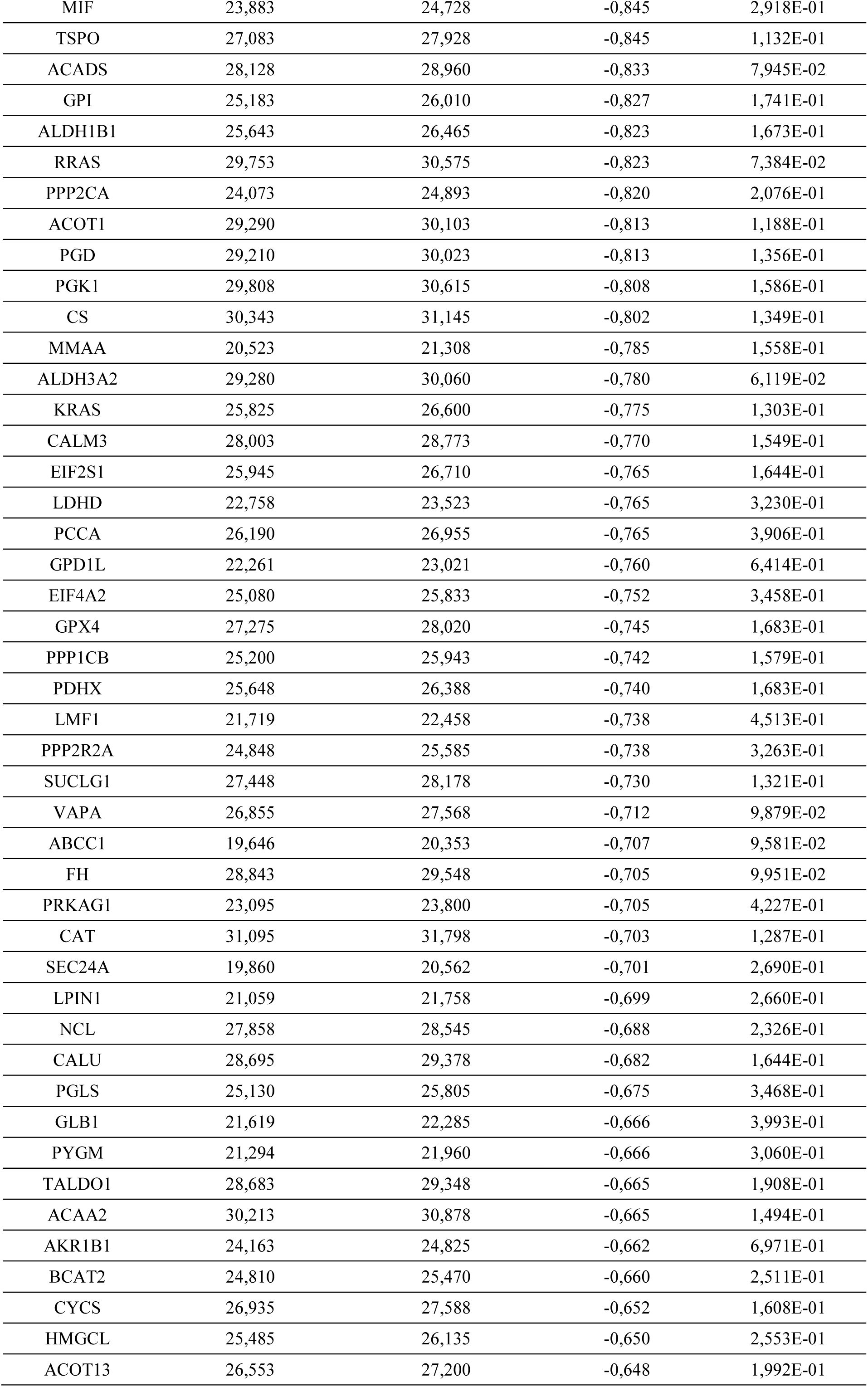
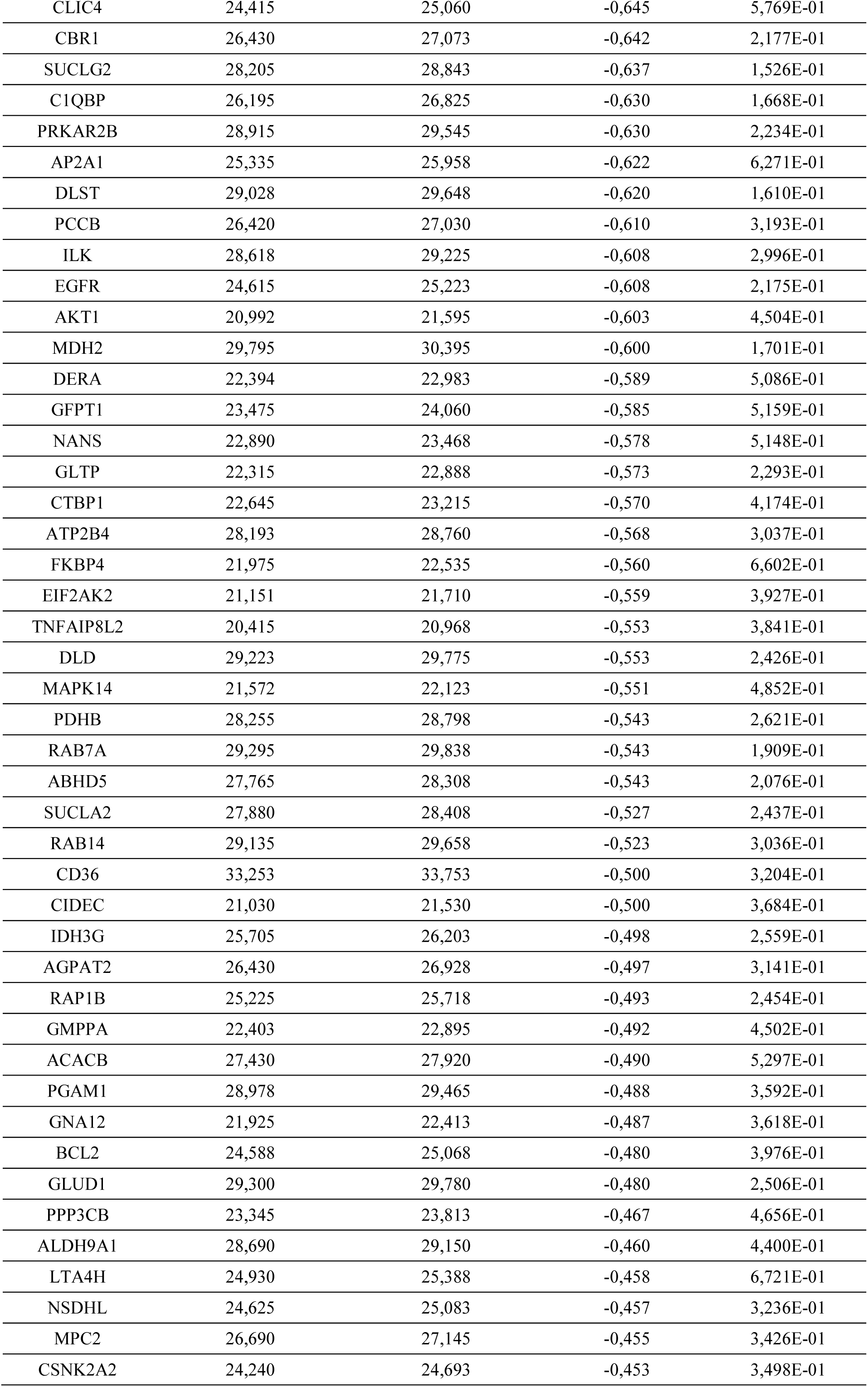
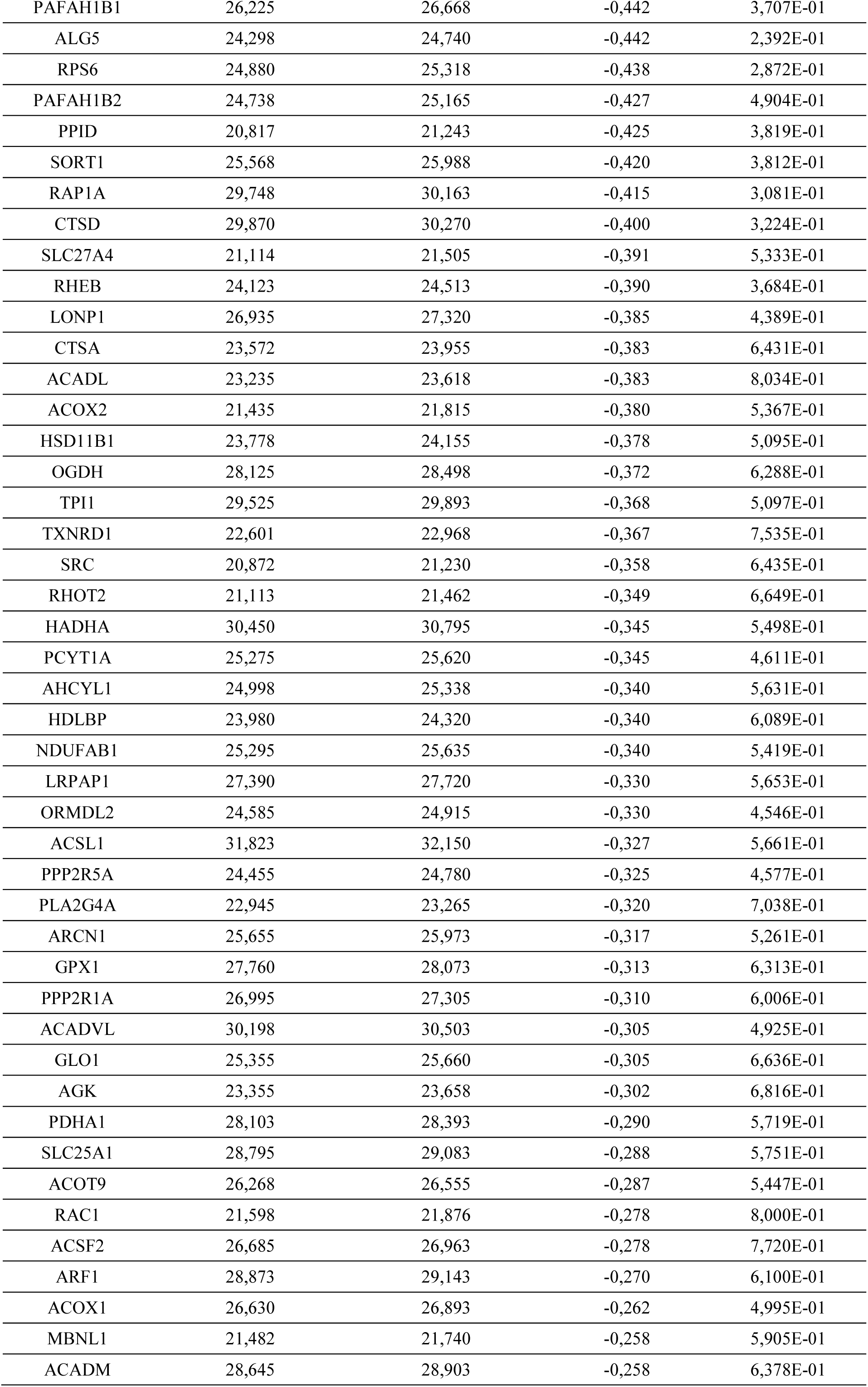
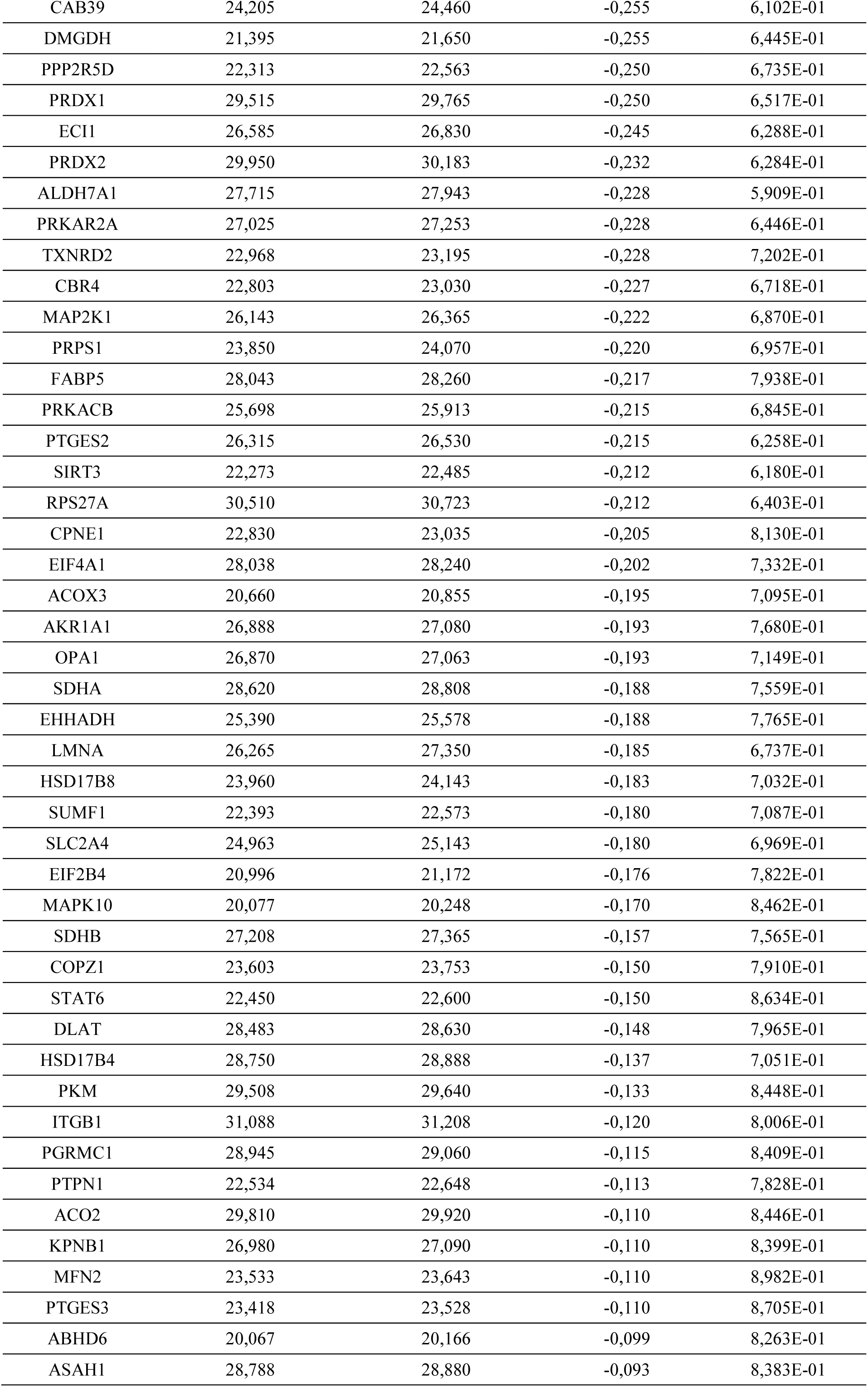
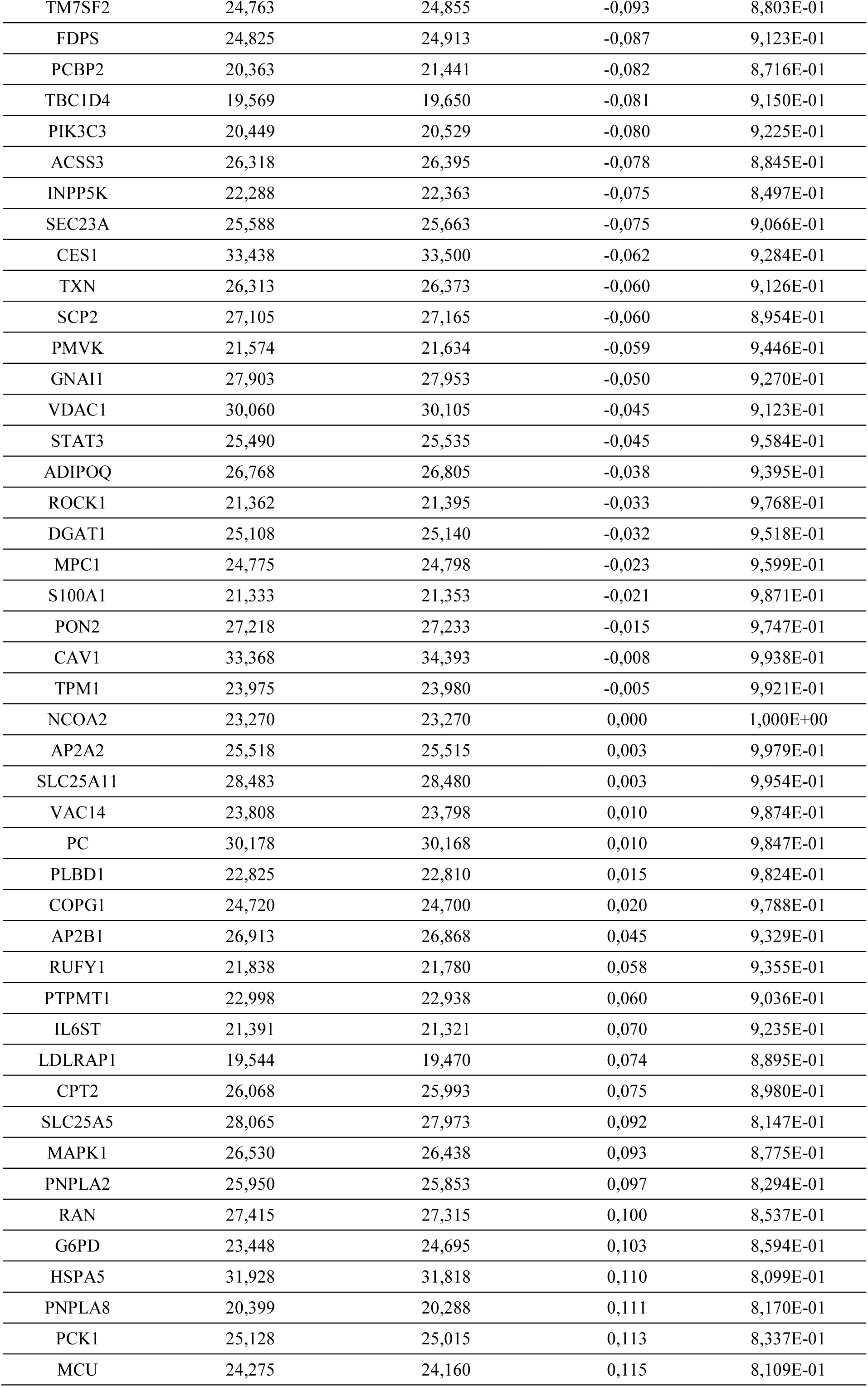
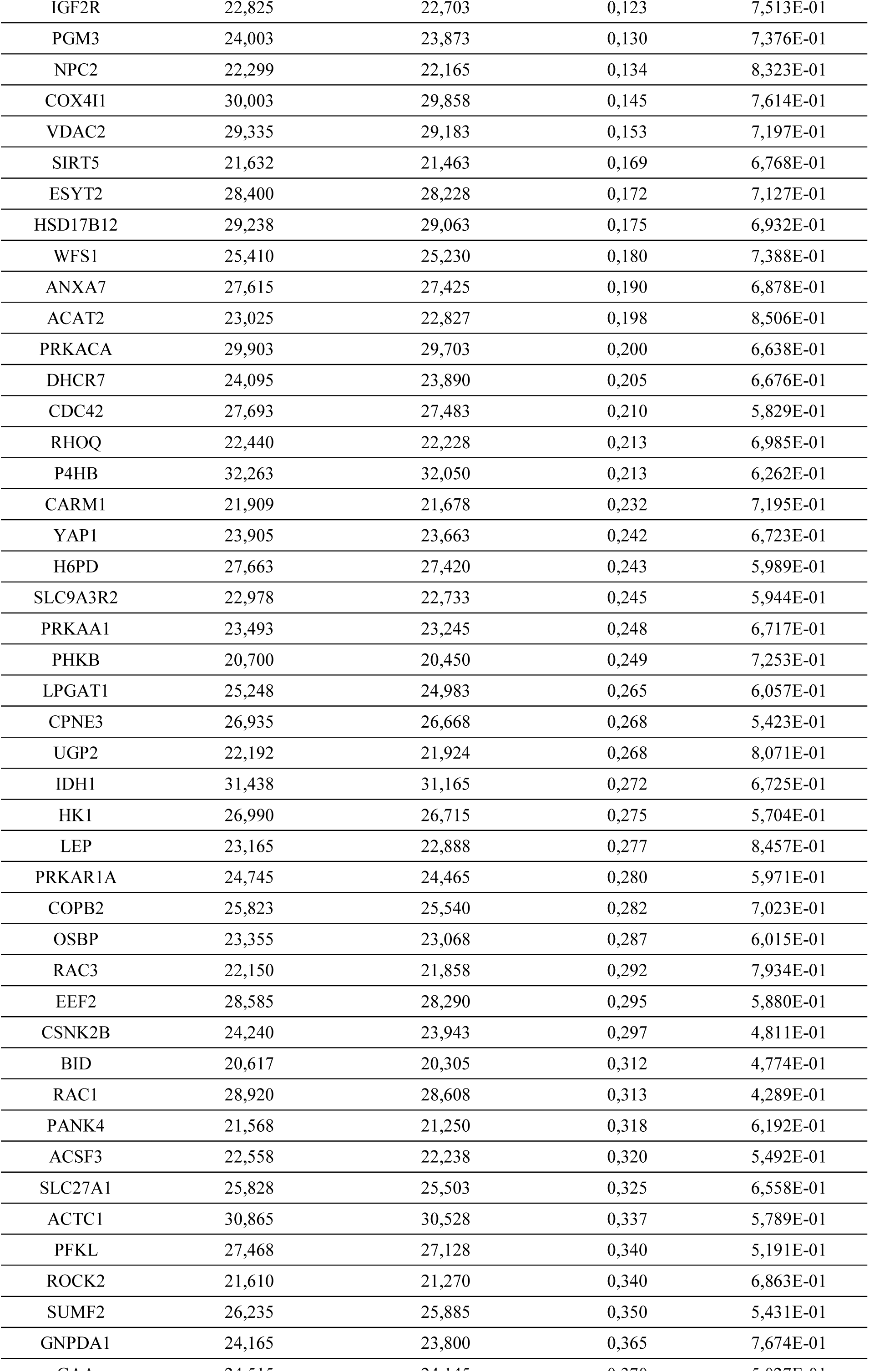
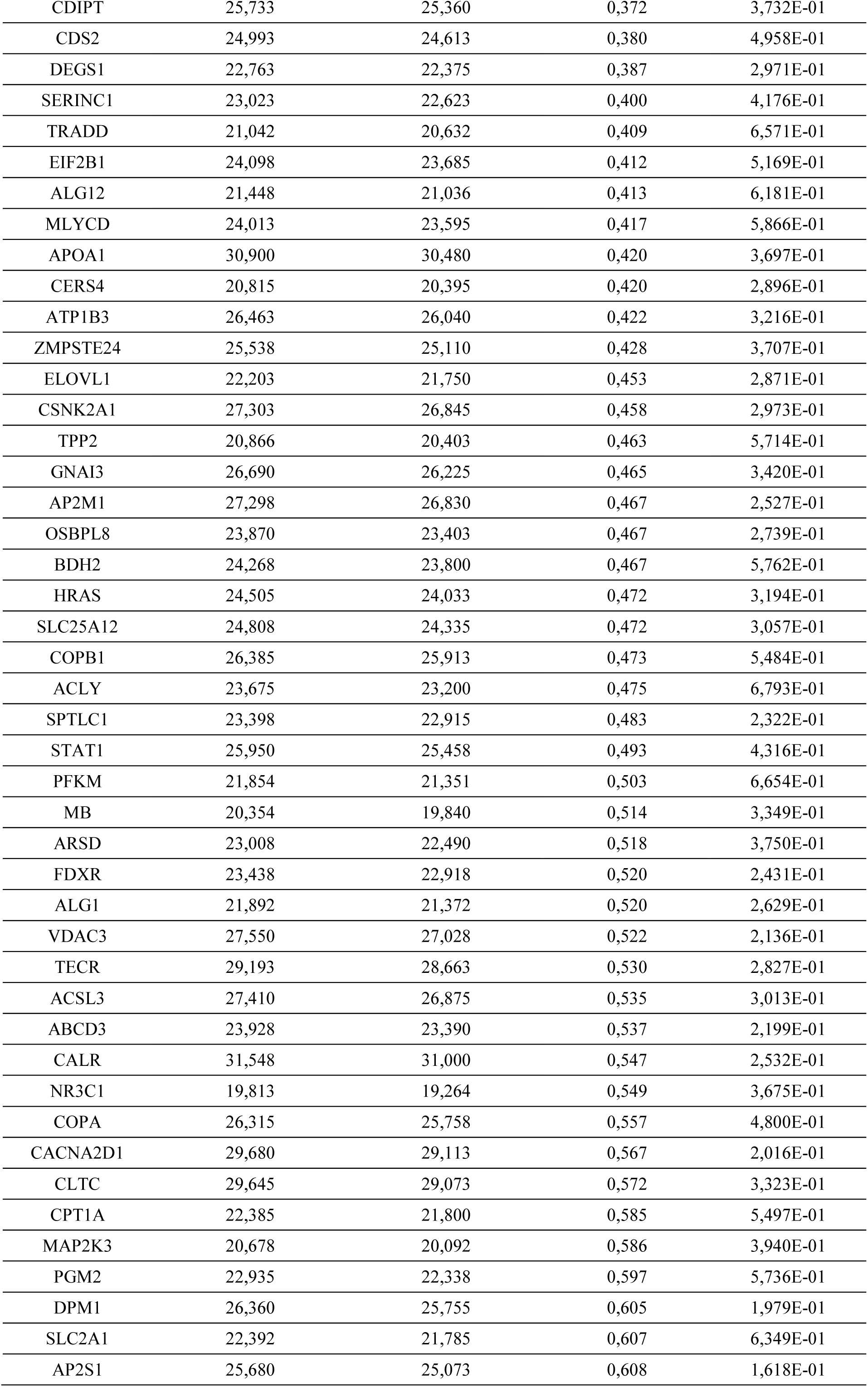
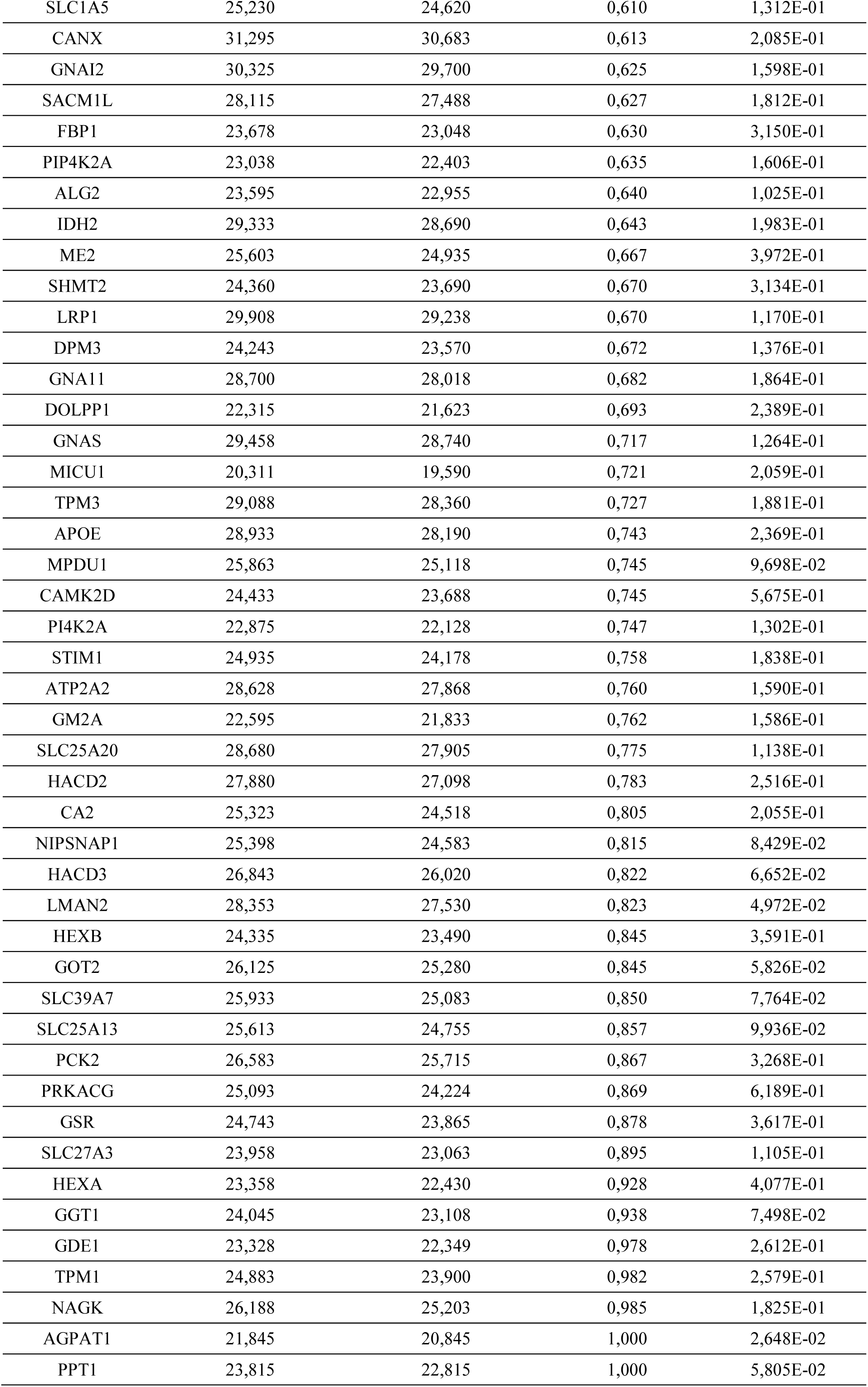
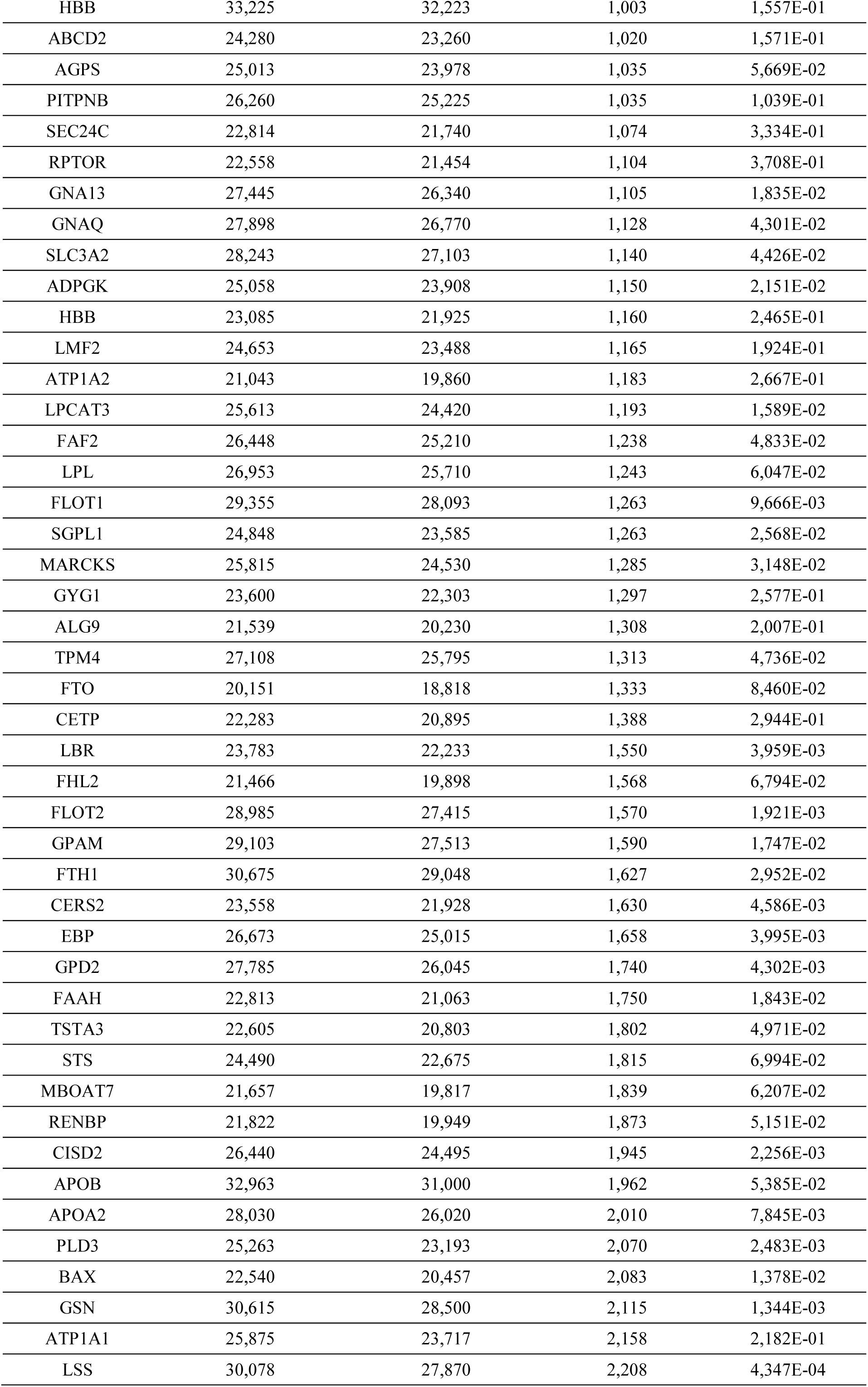
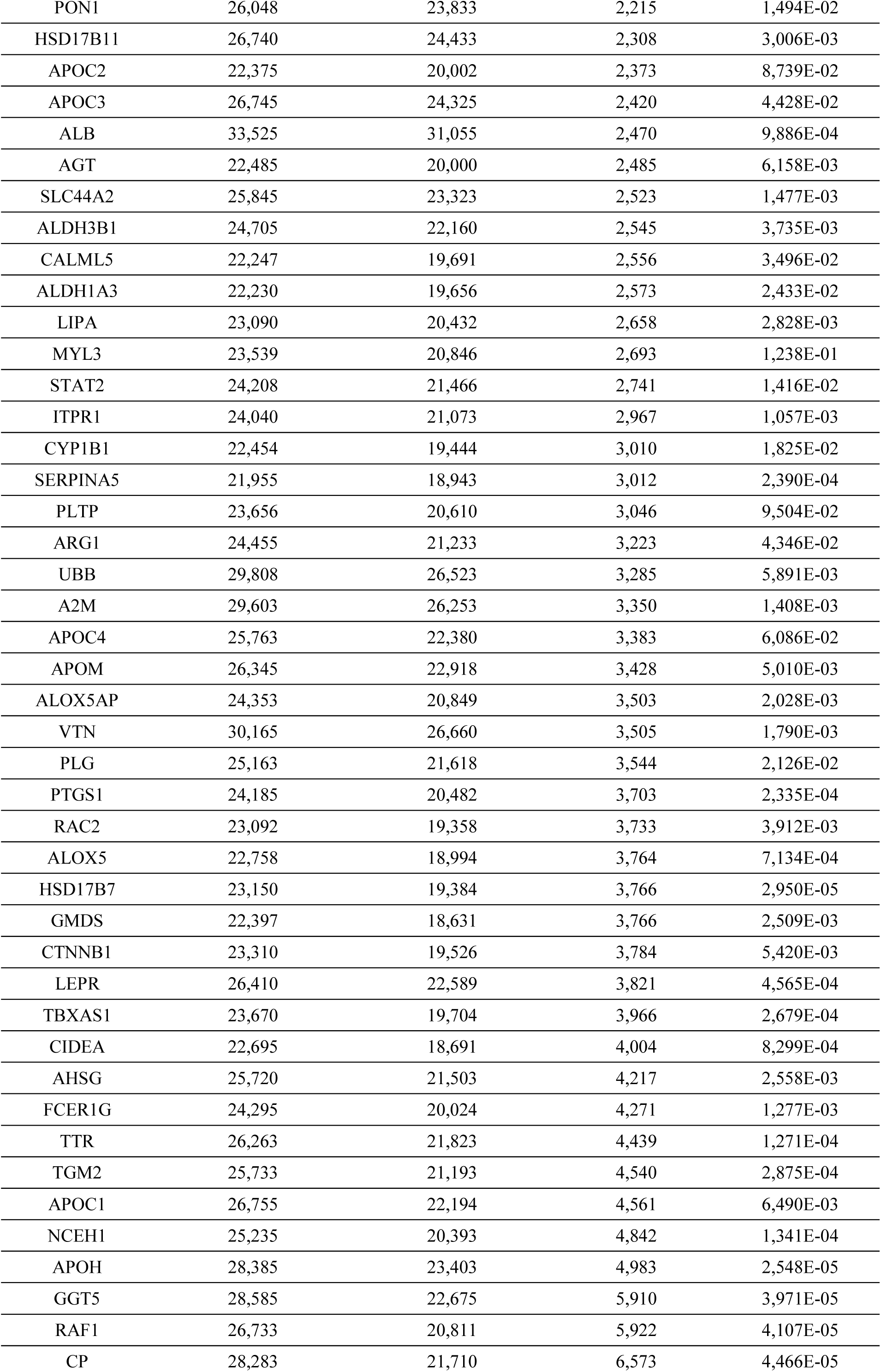
Protein expression involved in lipid and glucose metabolism were quantified by nano LC-MS/MS. The log2 transformed average intensities of label free quantification (LFQ) in BM-Ad and SC-Ad for each protein in the dataset and the corresponding log2 fold change and p-Value are presented.

